# FreeHi-C: high fidelity Hi-C data simulation for benchmarking and data augmentation

**DOI:** 10.1101/629923

**Authors:** Ye Zheng, Sündüz Keleş

**Affiliations:** Department of Statistics, University of Wisconsin – Madison, Madison, WI 53706, USA; Department of Biostatistics and Medical Informatics, University of Wisconsin – Madison, Madison, WI 53706, USA

## Abstract

Ability to simulate realistic high-throughput chromatin conformation (Hi-C) data is foundational for developing and benchmarking statistical and computational methods for Hi-C data analysis. We propose FreeHi-C, a data-driven Hi-C simulator for simulating and augmenting Hi-C datasets. FreeHi-C employs a non-parametric strategy for estimating interaction distribution of genome fragments from a given sample and simulates Hi-C reads from interacting fragments. Data from FreeHi-C exhibit higher fidelity to the biological Hi-C data compared with other tools in its class. FreeHi-C not only enables benchmarking a wide range of Hi-C analysis methods but also boosts the precision and power of differential chromatin interaction detection methods while preserving false discovery rate control through data augmentation.

## Introduction

Recent maturation of chromosome conformation capture (3C)^1^ and Hi-C sequencing technologies^2, 3^ led to high-throughput profiling of three-dimensional chromatin architecture and revealed transformative insights on long-range gene regulation^4–6^. Alongside the technological breakthroughs, a growing number of methodologies and algorithms^7–16^ emerged for the analysis of Hi-C and other 3C-derived data types. These methods are developed and benchmarked on disparate biological and simulated or computationally-constructed datasets that are often customized for the methods under the study. Furthermore, the high sequencing costs of Hi-C experiments, e.g., billions of reads for decent 10-40kb analysis resolution of mammalian genomes ^3, 17^, preclude the possibility of generating biological benchmarking datasets with sufficient sequencing depth, numbers of replicates, and with known true interactions.

Benchmarking and evaluation of Hi-C methods have relied on either conducting distance-stratified permutations of the contact matrices^15^ to nullify the underlying biological structure or simulating contact matrices directly by capturing only the most general spatial structures for specific analysis purposes^9, 11, 13, 14, 18, 19^. Additionally, pooling biological replicates and then randomly partitioning to generate pseudo-replicates and downsampling are also common approaches to study similarity metrics or evaluate the sequencing depth effect^12, 15, 18, 19^. A more systematic Hi-C simulation method, Sim3C^20^, has been recently proposed for the design of Hi-C experiments with respect to the selection of restriction enzyme, determination of sequencing depth, and power analysis. However, Sim3C imposes strong assumptions to parametrically mimic Hi-C contact matrix structures and arbitrarily introduces random domain structures, e.g., topological associating domains (TADs). The properties of actual chromatin interactions at different scales^11, 21–23^ are far more nuanced than the basic factors that Sim3C takes into account. Consequently, the resulting Sim3C simulated contact matrices do not visually resemble biological Hi-C contact matrices (Fig. 1a).

**Figure 1.**
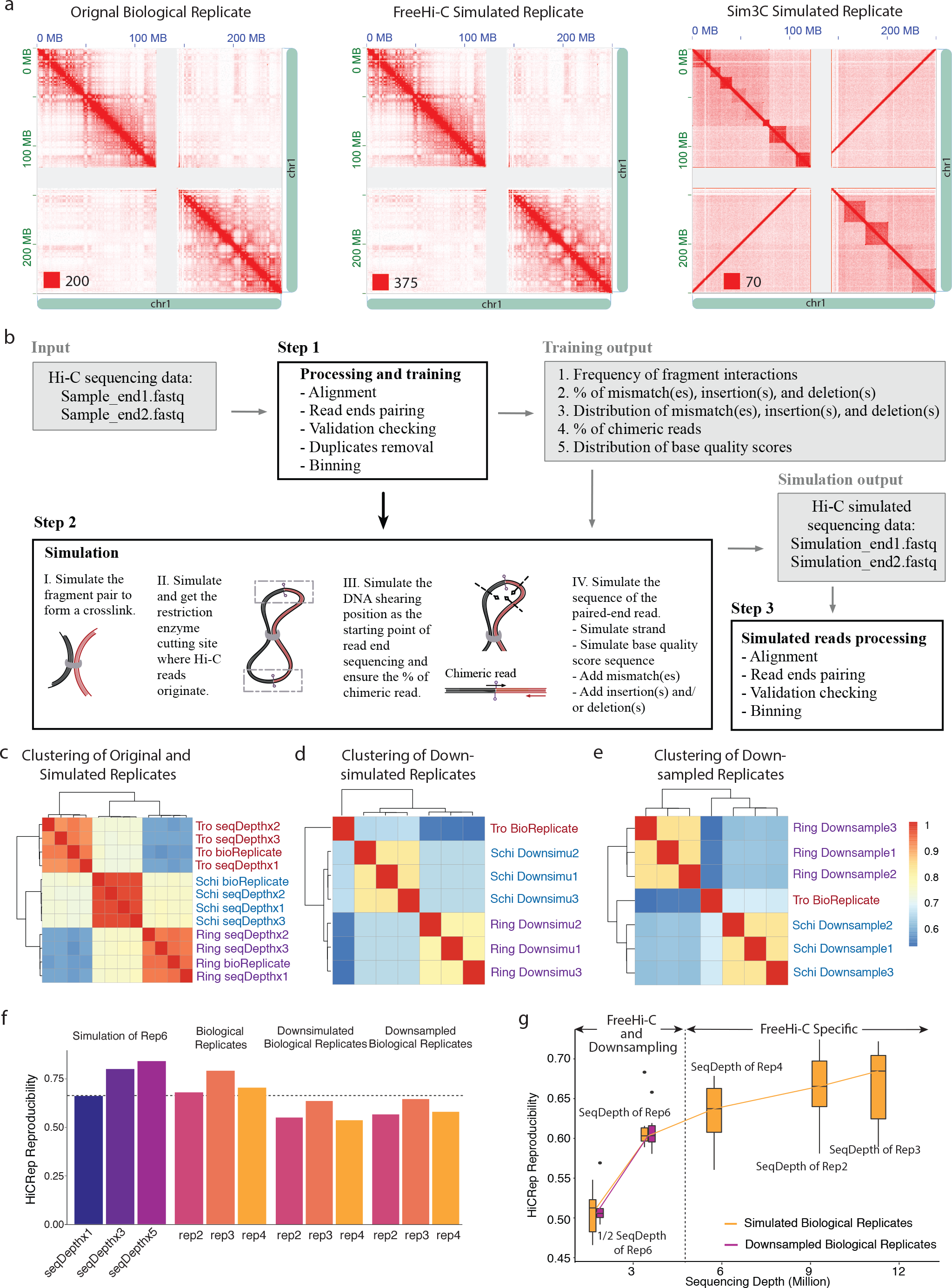
FreeHi-C enables simulating high fidelity Hi-C data. **a.** Hi-C contact matrix of biological rep2 of GM12878 along with FreeHi-C and Sim3C^20^ simulated replicates at matching sequencing depths. FreeHi-C and Sim3C parameters are set based on rep2. The numbers at the left bottom of the matrices are the color scale for the interaction counts. **b.** Details of the FreeHi-C simulation workflow. **c.** Hierarchical clustering of the original Hi-C biological replicates and the FreeHi-C simulated replicates for the Rings, Trophozoites, and Schizonts of *P. falciparum*. **d.** Hierarchical clustering of the FreeHi-C simulated replicates matching the sequencing depth of the original *P. falciparum* Trophozoites stage sample. **e.** Hierarchical clustering of the downsampled replicates matching the sequencing depth of the original *P. falciparum* Trophozoites stage. **f.** HiCRep reproducibility of the contact matrices between the biological replicate6 of GM12878 and corresponding FreeHi-C simulated replicates of different sequencing depths (first panel) as well as other biological replicates (second panel), FreeHi-C simulated biological replicates (third panel) or downsampled biological replicates (fourth panel) matching the sequencing depth of replicate6. **g.** Hi-CRep reproducibility of the contact matrices between pairs of biological replicates of GM12878 simulated by FreeHi-C (orange boxes and lines) or downsampled (purple boxes and lines) to 1/2 × sequencing depth of replicate6, 1 × sequencing depth of replicate6, and sequencing depths of replicate4, replicate2, and then replicate3, respectively.

## Results

In this paper, we present a robust Hi-C simulation method, FreeHi-C (**Fr**agment interactions **e**mpirical **e**stimation for fast simulation of **Hi-C** data) that simulates realistic read-level Hi-C data leveraging a nonparametric framework. FreeHi-C takes as input raw Hi-C sequencing reads in FASTQ format and estimates the frequency of genomic fragment interactions. This is fundamentally different from existing methods that simulate Hi-C contact matrices under a series of assumptions^9, 11, 14, 15^. Subsequently, FreeHi-C generates pairs of sequencing reads that represent the interacting fragment pairs with embedded random nucleotide mutations and indels while conserving the proportion of chimeric reads. Thereby, the variability and read-level characteristics of original Hi-C sequencing libraries can be preserved in the simulated sequences. Overall, the entire simulation procedure of FreeHi-C emulates the standard Hi-C experimental protocol^2, 3^ (Fig. 1b) where each read pair is generated independently, thereby allowing parallelization for faster implementation and the ability to simulate to the user-specified sequencing depth. The ability to simulate larger depths enables performance assessment of Hi-C analysis methods such as significant interaction and differential interaction detection with respect to varying sequencing depths. The method can be used to simulate Hi-C data from any organism, with any read length, and restriction enzymes. FreeHi-C is freely available at https://github.com/keleslab/freehic and includes a complete standard Hi-C data processing module ^8, 24^ encompassing raw sequence alignment, validation checking, duplicate removal, and genome binning. Hence the output is compatible with the standard input of downstream analysis such as normalization. Simulation component is an order of magnitude less time consuming than processing of the sequencing reads (Supplementary Fig. 1).

We illustrated the versatile features of FreeHi-C with Hi-C datasets of two human cell lines, GM12878 and A549, and malaria parasite *Plasmodium falciparum* 3D7, as representatives of large and small genomes, respectively. Analysis of GM12878 and A549 are carried out at 40kb resolution and 10kb for *Plasmodium falciparum* 3D7. Replicates are simulated at the same sequencing depth as the original biological replicates unless explicitly stated. For illustration purposes, we represent the analysis results utilizing all the interactions involving chromosome 1 of biological replicates, rep2, rep3, rep4, and rep6, for GM12878 and rep1 to rep4 for A549 cell line and all the 14 chromosomes from three life stages of *P. falciparum*. The sequencing depths of the replicates are summarized in Supplementary Fig. 2.

We first assessed how well FreeHi-C simulated replicates capture the general data characteristics of the biological replicates. Juicebox^25^ visualizations of the contact matrices of a FreeHi-C simulated dataset compared to the seed biological replicate and Sim3C realization with matching parameters are displayed in Fig. 1a. FreeHi-C simulated contact matrix exhibits markedly higher fidelity to the original biological replicate. Furthermore, the local interaction visualization in Supplementary Figs. 3-5 illustrates that FreeHi-C robustly captures the detailed chromatin interaction structures, such as the chromatin loops and TADs, for all biological replicates at any genomic location. Next, comparison of the distribution of contact counts stratified by the genomic distance between the simulated and biological replicates shows that simulated reads of the original sequencing depth give rise to contact count distributions similar to those of the seed biological replicates (Supplementary Figs. 6 and 7). After establishing high fidelity of individual simulation replicates, we assessed whether the simulated data of different biological replicates preserved the relationships existing between biological replicates. Fig. 1c displays hierarchical clustering of three biological *P. falciparium* samples together with three FreeHi-C simulated replicates each. The high fidelity of simulations visualized in Supplementary Fig. 8 leads to each simulated replicate being clustered with its seed biological sample and, furthermore, retains the relationship that chromatin architecture in Rings stage is more similar to that of Schizonts stage while Trophozoites stage has striking differences from the other two^26^. In addition to recapitulating the clustering structure of the biological samples, when compared in pairs, simulated replicates also display similar distance stratified log-fold-change trends of the contact counts as the biological seed samples (Supplementary Figs. 9 and 10).

Next, we demonstrate how FreeHi-C enables benchmarking a wide range of Hi-C analysis methods and compare it with the commonly used downsampling approach. Our first analysis focuses on the assessment of the reproducibility of Hi-C contact matrices by HiCRep^12^. Although Fig. 1c successfully clustered Rings and Schizonts stages together based on the HiCRep reproducibility quantification, this result can be challenged as biased due to the significant differences in the sequencing depths of the samples, i.e., Rings and Schizonts stages have 45-375% more reads (Supplementary Fig. 2). Therefore, it is desirable to adjust for the sequencing depth either by simulation or downsampling and re-evaluate the performance of HiCRep. Fig. 1d and Supplementary Fig. 11 illustrate that by down-simulating Rings and Schizonts stages to the smallest sequencing depth sample Trophozoites or up-simulating Trophozoites and Rings stages to that of the largest sample Schizonts, HiCRep reproducibility can consistently cluster Rings and Schizonts stages together. Downsampling, however, leads to Schizonts being mis-clustered with Trophozoites (Fig. 1e) indicating that downsampling may generate low-quality Hi-C matrices (i.e., reproducibility of Hi-C matrices from downsampled datasets ranged between 0.32 and 0.91) that potentially lead to wrong inference. To further explore the impact of removing sequencing depth bias on HiCRep reproducibility, we applied HiCRep between biological and the corresponding simulated replicates of GM12878 data. Supplementary Fig. 12a shows that, as expected, the HiCRep quantified reproducibility of the simulated replicates increases with the growing sequencing depths. However, we observe that the deeply sequenced biological replicates, e.g., rep2 and rep3 of GM12878, can achieve as high reproducibility with the smaller biological replicates as the low sequenced simulated replicates of rep4 and rep6 (Supplementary Fig. 12a). This contradicts the expectation that reproducibility of a biological replicate with another independent biological replicate cannot exceed the reproducibility with its FreeHi-C simulated replicate. By simulating the biological replicates, for example, rep2, rep3, and rep6, to the sequencing depth of the target sample rep4, we eliminate the sequencing depth bias and unveil the underlying similarity relationship (Supplementary Fig. 12b). In contrast, downsampling strategy can only be implemented for the smallest replicate, namely downsampling rep2 to rep4 to sequencing depth of rep6 (Fig. 1f). Hence, FreeHi-C facilitates evaluation of HiCRep and similar methods at a wider range of sequencing depths without any limitation (Fig. 1g). We provide a similar sequencing depth debiasing discussion for Fit-Hi-C^7^ in detecting significant interactions in Supplementary Materials.

Another pivotal benchmarking utility of FreeHi-C is for comparing performances of different methods that address the same Hi-C inference problem. We considered the detection of differential chromatin interactions (DCIs) as an example and compared the state-of-the-art differential interaction identification methods, diffHic^9^ and multiHiCcompare^16^, across a series of sequencing depths. We singled out these two approaches because alternatives such as FIND^13^ and HiCcompare^14^ have either established lack of false discovery rate control and excessively long run times^14, 19^ or constituted a specialized version of multiHiCcompare to exclusively handle one replicate per condition. We compared diffHic and multiHiCcompare with respect to false discovery rate (FDR) control and power through both FreeHi-C simulation and down-sampling. Overall, Supplementary Fig. 13 elucidates that both methods can control the FDR when replicates are evaluated for DCIs within the GM12878 data. Remarkably, downsampling displays an increasing FDR trend as the sequencing depth increases; however, this trend can only be investigated up until the original depth of the samples since the sequencing depths of downsampled samples cannot go beyond their original depths. FreeHi-C simulation elucidates the trends when the depths of the samples go beyond their original ones and illustrates conservative FDR control at higher depths. This highlights a clear pitfall of drawing conclusions solely based on downsampling. Additionally, downsampling and FreeHi-C simulation have distinct implications for detection power of the methods (the first column of Supplementary Fig. 14). FreeHi-C consistently indicates higher detection power for multiHiCcompare than diffHic; however, downsampling reverses the rank of the two methods and only reveals the same conclusion as FreeHi-C after small log-fold-change interactions are filtered (the second and third columns of Supplementary Fig. 14). Collectively, FreeHi-C assesses performances of competing methods without any sequencing depth limit. Downsampling, however, can only accommodate comparisons at lower sequencing depths of the available samples and may lead to inconsistent or unreliable discoveries.

A key impediment for differential chromatin interaction inference with Hi-C data is the limited numbers of biological replicates. As a result, numerous practical challenges arise. We identified three commonly encountered scenarios as a function of the number of replicates available per condition (Fig. 2) and assessed how FreeHi-C simulated replicates can offer advancements. These three settings are one replicate per condition (ORPC), uneven numbers of replicates per condition where one of the conditions have only a single replicate (URPC), and multiple numbers of replicates per condition (MRPC). Under these settings, we applied multiHiCcompare which demonstrated stronger power in the previous analysis, for differential chromatin interaction detection. For all the settings, we first provide the assessment that established false discovery rate control and then investigate detection power and the precision of top significant findings, where the set of differential interactions from the full set of biological replicates (4 from each of the A549 and GM12878 cells) is utilized as the gold standard. In this first setting ORPC where each condition has only one replicate, the false discovery rate control with the BH procedure^27^ ensures that the observed FDR is below the target FDR (the first panel in Fig. 2a and Supplementary Fig. 15); however, overall, this setting exhibits extremely conservative FDR control. This comes, not so unexpectedly, at the cost of low power (the third panel in Supplementary Fig. 16). When we augment each of the conditions with FreeHi-C simulated replicates, FDR is still well controlled (Fig. 2a and Supplementary Fig. 15), and power increases by an average of 300 folds across the five levels of FDR thresholds, going from only a few hundred differential interactions to a few tens of thousands (Supplementary Fig. 16b). It is reasonable to argue that under ORPC, the key target should not be a discrete, thresholded list of DCIs. Instead, the focus should be on the ranking of the interactions because the top significant DCIs are typically utilized for downstream analysis and experimental follow-up. We scrutinized the accuracy of the top significant DCIs identified with and without FreeHi-C augmentation by comparing the ranked lists to the set of DCIs identified by utilizing all the 4 biological replicates of the two conditions. Fig. 2b illustrates that FreeHi-C simulated samples lead to a significantly higher precision of 100-75% compared to ~10% of the ORPC analysis (evaluations under additional gold standard settings are provided in Supplementary Fig. 17). We next evaluated the biological relevance of the “ranked up” and “ranked down” DCIs due to FreeHi-C augmentation by external RNA-seq and CTCF ChIP-seq data. The new ranking of the top DCIs overlapped significantly more with the differentially expressed genes of the two cell lines (Supplementary Fig. 18a), supporting FreeHiC simulation augmentation. Overlap with differential CTCF ChIP-seq peaks deemed both the original and the new rankings similarly significant (Supplementary Fig. 18b) except for a clear advantage for ranking with the FreeHiC augmentation at the very top 100 DCIs. An immediate extension of the ORPC includes uneven numbers of replicates per condition, particularly, single replicate for one and two or three replicates for the other. URPC may arise either by design or due to low quality or failed replicates. Figs. 2c and d for URPC highlight that, similar to ORPC, FreeHi-C augmentation of the conditions markedly boosts power (Supplementary Fig. 19b), refines ranking of the DCIs (Supplementary Figs. 20-22) while preserving FDR control (Supplementary Fig. 19a), and exhibiting strong genomic support (Supplementary Fig. 23).

**Figure 2.**
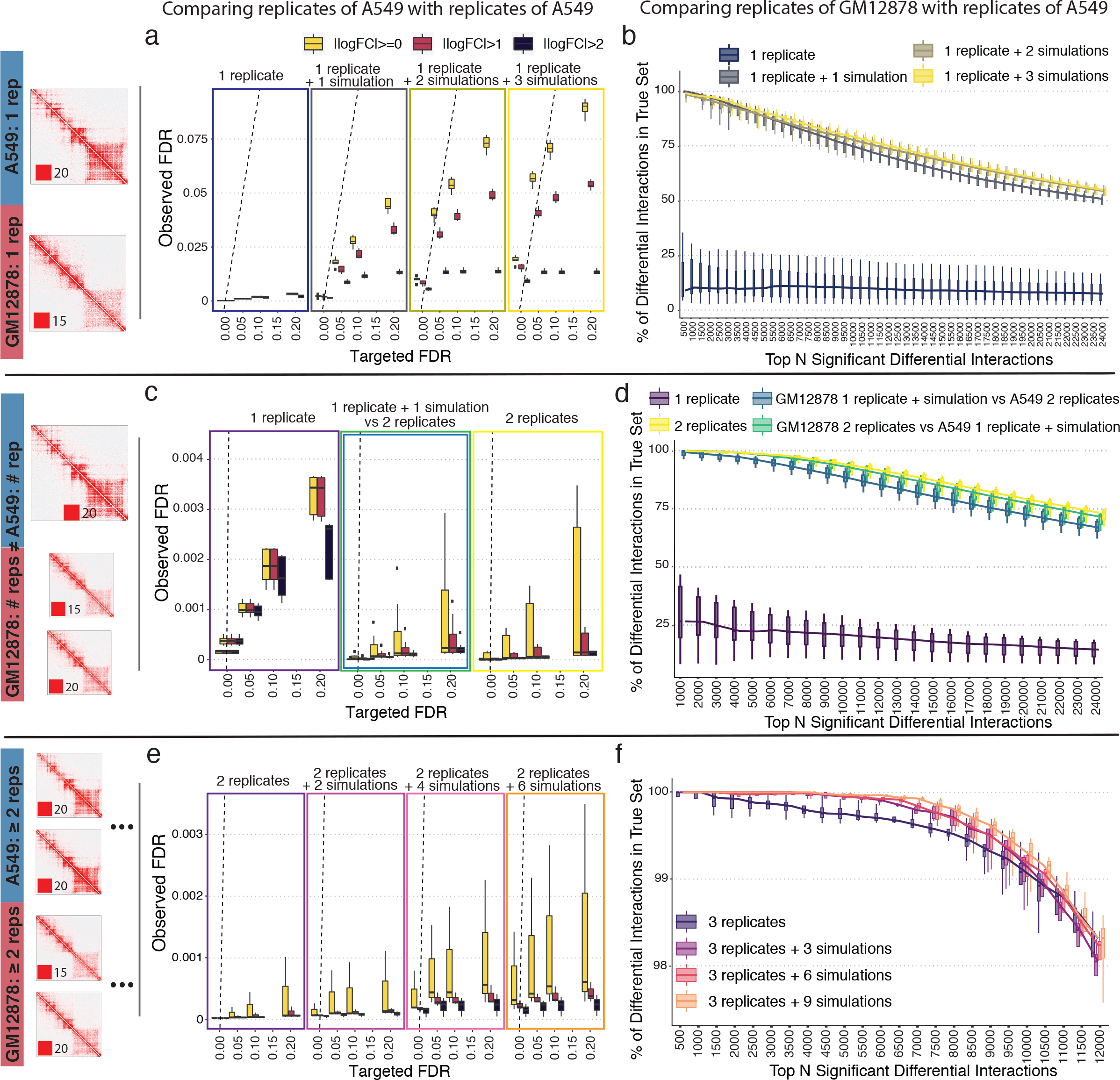
Data augmentation with FreeHi-C simulated replicates improves differential chromatin interactions (DCIs) detection. Rows 1 to 3 refer to one replicate per condition (ORPC), uneven numbers of replicates per condition (URPC), and multiple replicates per condition (MRPC) settings, respectively. **a**, **c**, and **e** delineate observed false discovery rates of within-sample comparisons for A549 data (i.e., comparisons of replicate(s) of A549 with other replicate(s) of A549). The dashed lines are y = x. **b**, **d**, and **f** display precision, computed as the percentage of top significant DCIs of each specific analysis in the “true” differential chromatin interaction list, as a function of top-ranking DCIs. The true set is defined by comparing the full set of 4 replicates of GM12878 with 4 replicates of A549.

Finally, we generalized the FreeHiC augmentation strategy as a meta-analysis approach for the general setting of multiple replicates per condition (Figs. 2e and f). To account for the fact that simulated samples cannot provide additional full degrees of freedom, the meta-analysis approach pairs biological and simulated samples in numbers concordant with the original differential testing design and aggregates p-values of candidate differential interactions across comparisons by Fisher’s method^28^. It then refines the significance ranking for FDR control with the BH procedure by taking the median of the adjusted p-values per interaction (Methods). Fig. 2e and Supplementary Fig. 24 reveal that FDR control is well preserved as the number of simulated replicates in augmentation increases. Notably, augmentation with simulated replicates not only boosts the number of significant DCIs identified across a series of FDR thresholds (Supplementary Fig. 25), but also results in a significantly better ranking with higher precision of the top significant DCIs for further quantitative and experimental validation (Fig. 2f and Supplementary Figs. 26-31). We further evaluated the biological relevance of the DCIs identified with meta-analysis via FreeHiC augmentation by leveraging RNA-seq and CTCF ChIP-seq data. With the enrichment levels using the full set of 4 replicates per condition serving as the best achievable outcome (panel b of Supplementary Figs. 32 and 33), meta-analysis via FreeHi-C augmentation successfully improves the significance of the overlapping between DCIs and the differentially expressed genes (Supplementary Figs. 32 and 33 for MRPC with 3 and 2 replicates per condition, respectively). Evaluation with differential CTCF ChIP-seq peaks results in significant co-localization for all the detection settings with or without FreeHiC augmentation (Supplementary Fig. 34).

Analytical methods for analyzing data from Hi-C and related experiments are growing at a faster pace than they are benchmarked and evaluated uniformly. We developed FreeHi-C to enable simulating read-level Hi-C data in a data-driven manner. A direct advantage of FreeHi-C non-parametric construction by closely following the Hi-C experimental protocol is that contact matrices resulting from FreeHi-C simulated reads accurately capture the nuanced interaction structures of real Hi-C contact matrices. We showcased the benchmarking utility of FreeHi-C simulations by investigating the performance of HiCRep^12^ for reproducibility quantification, Fit-Hi-C^7^ for significant interaction detection, and diffHic^9^ and multiHiCcompare^16^ for identifying DCIs. We demonstrated the outperformance of FreeHi-C over the downsampling approach with respect to flexibility and reliability as a benchmarking tool. Most notably, we illustrated that data augmentation with FreeHi-C simulation improves detection of differential chromatin interactions across all the settings parametrized by the numbers of available replicates per condition while preserving FDR control. FreeHi-C augmentation led to markedly raises the accuracy of ranked DCIs with better biological support compared to analysis only with the original replicates. The robust performance of FreeHi-C data augmentation leads to more accurate lists of DCIs which are essential for effective downstream analysis.

## Acknowledgments

This work was supported by NIH grants HG009744 and HG007019 to Sündüz Keleş.

## Author Contributions

S.K. and Y.Z. conceived the project. Y.Z. and S.K. designed the research and developed the method. Y.Z. developed the simulation framework and performed the experiments. Both authors contributed to the preparation of the manuscript.

## Competing Interests

The authors declare no competing financial interests.

## Methods

### FreeHi-C simulation framework

#### Processing and training module

FreeHi-C implements the steps outlined in Fig. 1b. It takes as input raw Hi-C sequencing data in the form of FASTQ files, processes the reads, and learns the parameters for the simulation module. The sequence processing module follows a standard protocol^24^ by aligning raw paired-end read files individually and then joining the read ends to form read pairs, followed by interaction validation checking and duplicate removal steps. After obtaining the valid read-pairs (i.e., interactions), it fits an interaction-level mixture model to estimate the genomic fragment interaction frequencies, with the genomic fragments defined by the experimental restriction enzyme cutting sites. This sampling model considers two multinomial distributions with parameters 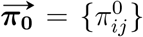 and 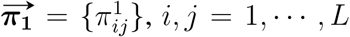, where *L* denotes total number of genomic fragments, for generating interaction events. The distribution indexed by the parameter *π*_0_ reflects background interactions, driven by experimental artifacts such as genomic distance whereas the second distribution, *π*_1_, characterizes true biological interaction signals. For each genome fragment “interaction” event, one interaction is drawn from each of the multinomial distributions, and one of them is recorded as the actual interaction event with probability *α*. Consequently, the distribution specified by this interaction-level sampling model is multinomial with vector-valued parameter 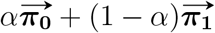. This intuitive sampling model, estimation of which does not require deconvolution of the two distributions, is key for FreeHi-C’s successful capture of the interactions in a given biological sample. Once the genomic fragment interaction is sampled, FreeHi-C generates a read pair from this interaction by taking into account strand configuration of the ends of the read pair, mismatch(es), insertion(s), deletion(s), the proportion of chimeric reads, and base quality scores of the reads. Hence, as part of the training module, FreeHi-C estimates these set of parameters empirically from the collection of valid reads. Specifically, FreeHi-C estimates the frequency distributions of numbers of insertions, deletions, and different types of mismatches across all the valid read pairs. Additionally, FreeHi-C processing and training module records the proportion of chimeric reads, i.e., reads pairs where one or both of the read ends are sequenced over the ligation junction, that are rescuable with the aim of preserving this proportion for the simulated reads. Finally, FreeHi-C empirically estimates the distribution of base quality scores for each locus of the Hi-C reads and uses these estimates to ensure that the simulated reads have similar base quality scores as the seed biological replicate.

#### Simulation module

FreeHi-C simulates fragment pairs from the estimated interaction-level mixture model as genomic fragments that form crosslinks in the Hi-C experiment protocol^3^. Ligation procedure in the experiment leads to two fragment junction sites. FreeHi-C randomly generates one of these which is then passed onto the next step to emulate DNA shearing. Two DNA shearing loci are randomly selected within ± 500bp, by default, of the selected ligation site. These two shearing loci also work as the starting points of the sequencing procedure. FreeHi-C extracts the sequences of the given length, for example, 36bp for 36bp paired-end sequencing, from these loci and assigns strand direction accordingly. During this sequencing step, reads closer than the requested read length to the ligation sites can be generated, as an emulation of chimeric reads. The final step is to introduce noise to the read sequences so that the mismatches and indels match to those in the reads of the original biological sample. Utilizing the empirical distribution of the sequence base quality scores across individual locus, FreeHi-C simulates such scores for each read at the nucleotide level. A key strength of FreeHi-C is that it can generate as many reads as specified by the user and outputs these in the FASTQ format. Furthermore, it processes the resulting reads according to the standard analysis protocol of Hi-C reads by the processing module. Through the post-simulation processing, FreeHi-C can directly provide genomics contact counts in a sparse matrix format (BED) compatible with the standard input format of downstream Hi-C analysis. Processing of the raw reads and learning of the parameters can be implemented on individual read pairs followed by a final aggregation; hence this module can be efficiently parallelized. Furthermore, simulations based on the same parameter settings are parallelized at the read-pair generation level.

### Simulation parameter settings

Different simulation parameter settings lead to variations in sequencing depth and also serve as a user guide for generating simulations that are more close or different than the seed biological replicate. Supplementary Fig. 35 considers a pre-processing parameter (resolution at 10kb or 40kb) and four different generative parameters (mutation and indel rates, utilization of chimeric reads, and sequencing depth) that impact the signal to noise ratio of the resulting Hi-C dataset. Further systematic evaluation of the reproducibility between simulation samples of varying depths and the seed biological sample emphasizes sequencing depth as the major factor that drives the similarity between simulated Hi-C contact matrix and the original one (Supplementary Fig. 36).

### Data augmentation with FreeHi-C simulated samples

When testing for differential chromatin interactions with two or more replicates per condition, we employed a FreeHi-C simulation augmented meta-analysis strategy. This approach generates simulation replicates for each of the original *n* biological replicates per condition and considers 2^*n*^−1 additional tests to preserve the degrees of freedom of the original test statistic. For example, for a setting with 2 biological replicates per condition, we generated 4 FreeHi-C simulation samples, one per original biological replicate, and evaluated the following comparisons, where *c*_1_ and *c*_2_ refer to the two testing conditions under consideration.

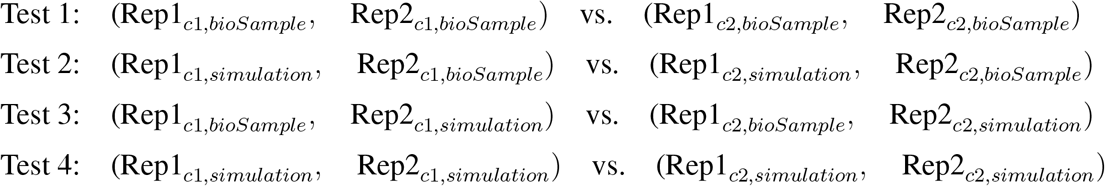

The p-values of the these tests are then aggregated by Fisher’s method^28^ as 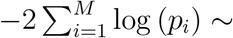 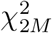, where *M* = 4 in the above example. Since the tests are not independent, the resulting aggregated p-values are anti-conservative. To dampen this effect, instead of ranking the differential interactions with the Fisher’s p-value in the BH procedure^27^, we rank them based on median of their adjusted p-values from individual test. Specifically, instead of ordering the hypothesis sequence *H*_(*i*)_, *i* = 1, 2, …, *M*, by the aggregated p-values obtained from Fisher’s method, we order them by the new significance rank determined by the median adjusted p-values of the individual tests and denote such ordering as *H*_(*r*(*i*))_, *i* = 1, 2, …, *M*, where *r*(*i*) is the index of the interaction that is ranked *i*^*th*^ in the new significant ranking list. Let *k* be the largest *i* for which 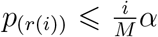, then the BH procedure rejects all *H*_(*r*(*i*))_, *i* = 1, 2, …, *k*. This is a more conservative procedure than the ordinary BH procedure on the Fisher’s p-values because *p*(*i*) ≤ *p*(*r*(*i*)); hence, the number of rejections, *k*, is always smaller or equal to the number of rejections using the BH procedure with Fisher’s p-values. The computational experiments support that this reverses the potential anti-conservative effect of aggregating dependent p-values with Fisher’s method.

### Evaluating the ranking of detected DCIs

The “true” differential interaction set is approximated by the most significant DCIs detected (FDR ≤ 0.001, 0.005, 0.01, 0.05, respectively) using 4 biological replicates of GM12878 versus 4 biological replicates of A549. In the sets presented in this paper, we rank the DCIs by their significance order and quantify the fraction of the top N significant differential interactions that appear in the “true” set where N varies as 500, 1000, etc. This quantity refers to recovery rate or precision.

Another type of “true” DCI list is specially defined for tests of uneven number of replicates per condition (Supplementary Figs. 21 and 22). In this setting, the true DCI set is defined as the set of most significant interactions in the comparison of rep2 and rep4 of GM12878 with rep1 and rep4 of A549. Accordingly, we measure the precision of the results from the following four comparisons: (i) one replicate out of rep2 and rep4 of GM12878 versus one out of rep1 and rep4 of A459; (ii) one replicate out of rep2 and rep4 of GM12878 with its FreeHi-C augmentation versus one out of rep1 and rep4 of A459 with its FreeHi-C augmentation; (iii) rep2 and rep4 of GM12878 versus one out of rep1 and rep4 of A459 with its FreeHi-C augmentation; (iv) one out of rep2 and rep4 of GM12878 with its FreeHi-C augmentation versus rep1 and rep4 of A459.

### Evaluating DCIs detected by FreeHi-C augmentation with RNA-seq and CTCF ChIP-seq

We evaluated the significance of the observed proportion of differentially expressed (DE) genes between GM12878 and A549 that overlap with significant DCIs using a randomization test. An empirical null distribution for the observed overlap statistics is constructed by randomly selecting an equal number of interactions as significant ones from all valid bin-pairs and overlapping these with the DE genes. The significance level of observed overlap is quantified by the percentage of random selection results that is larger than or equal to the observed statistics. A similar strategy is implemented for evaluating co-localization of DCIs with differential CTCF ChIP-seq peaks.

### Data availability

To study the operating features of FreeHi-C, we utilized two publicly available human Hi-C datasets as examples of large genomes with four replicates, rep2, rep3, rep4, rep6 from GM12878^3^ cell line and another four replicates, rep1-4, from A549^29^. Raw FASTQ files for GM12878 were downloaded from GEO^30^ under the accession GSE63525 and raw sequences for A549 were obtained from the ENCODE portal^31^ (https://www.encodeproject.org/) with accession ENCSR662QKG. For evaluation of FreeHi-C performance on small genomes, we leveraged three different stages of malaria parasite *Plasmodium falciparum* red blood cell cycles^26^. GM12878 and A549 are both processed at 40kb resolution, and *P. falciparum* at 10kb.

For validating the differential interaction detection with a differential expression analysis, we utilized RNA-seq gene expression data from the ENCODE portal (accession ENCSR000AED for GM12878 and ENCSR000CTM for A549). Similarly, the CTCF ChIP-seq peak signal files were also downloaded from ENCODE under accession ENCSR000DZN for GM12878 and ENCSR000DPF for A549.

### Software availability

FreeHi-C pipeline is implemented in Python with C accelerated core calculations and it naturally fits in the high-performance computing environments for parallelization. The source code and instructions for running FreeHi-C are publicly available at https://github.com/keleslab/FreeHiC.

## Supplementary Texts

In addition to studying the reproducibility of Hi-C datasets using HiCRep in a sequencing depth debiased manner, FreeHi-C simulations can also be leveraged to study sequencing depth dependent characteristics of Fit-Hi-C^7^ and similar methods in detecting significant interactions. A standard benchmarking approach for methods like Fit-Hi-C^7^ is to ask what proportion of the significant interactions are reproducibly recovered among different biological replicates. Similar to the findings of HiCRep reproducibility, the recovery ratio of the significant interactions is higher in the simulated replicates and substantial differences in sequencing depths can make the recovery rate of high depth biological replicates seem close to those of the lower depth simulated replicates of low depth biological replicates (Supplementary Fig. 37a). Simulation and downsampling of biological replicates to the sequencing depth of the target replicate enable debiasing of the comparison of significant interaction detection between simulated replicates and biological replicates from sequencing depth (Supplementary Fig. 37b and 38).

