## Supplementary Figures and Tables for "FreeHi-C: high fidelity Hi-C data simulation for benchmarking and data augmentation"

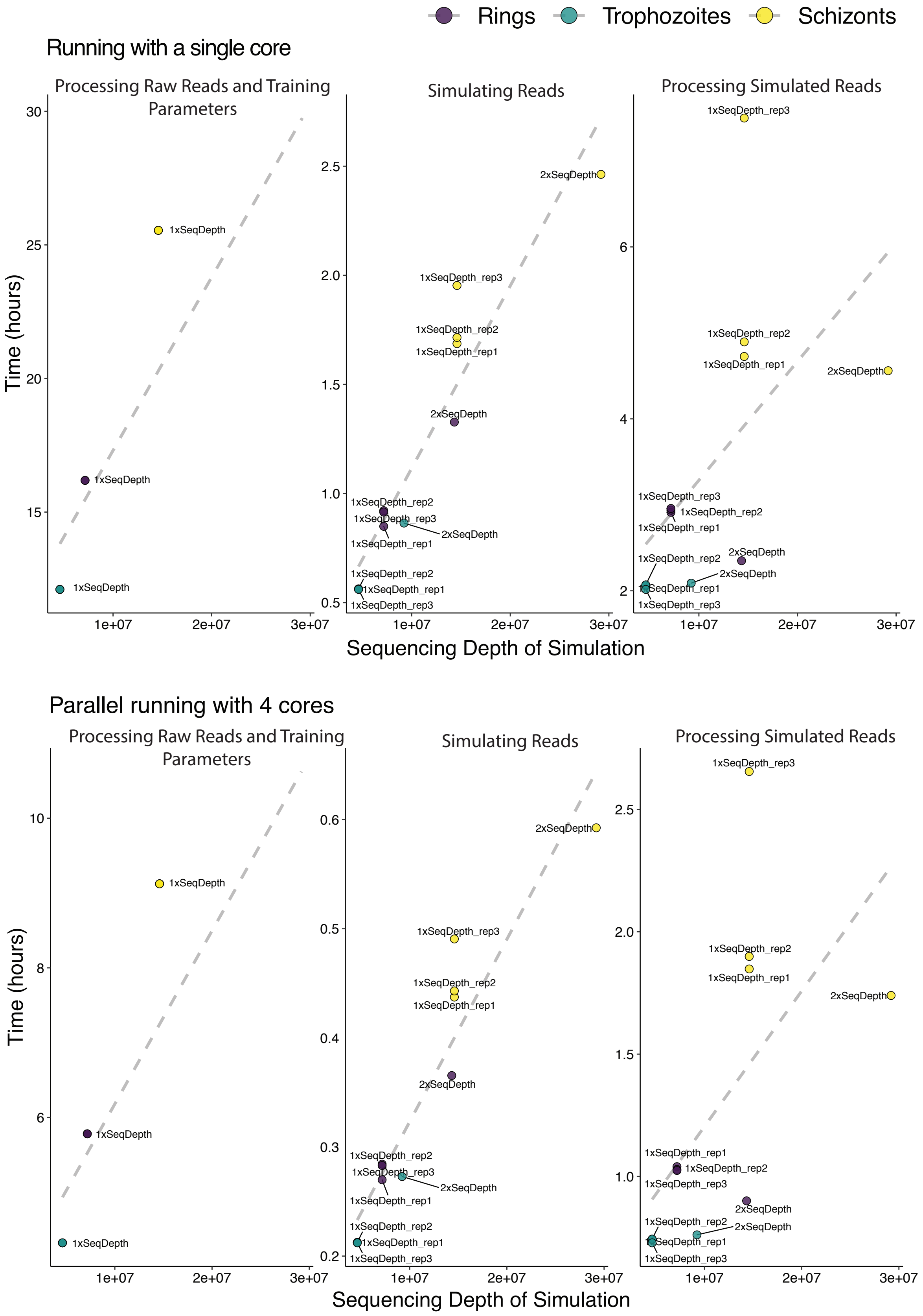

**Supplementary Figure 1 Runtime performance of FreeHi-C using a single core or four cores for parallel running.** Runtime quantification of three main steps of FreeHi-C workflow: processing of raw reads and parameter training, generating simulated reads, and processing simulated reads. Reported results are from *P. falciparum* Hi-C data simulations.

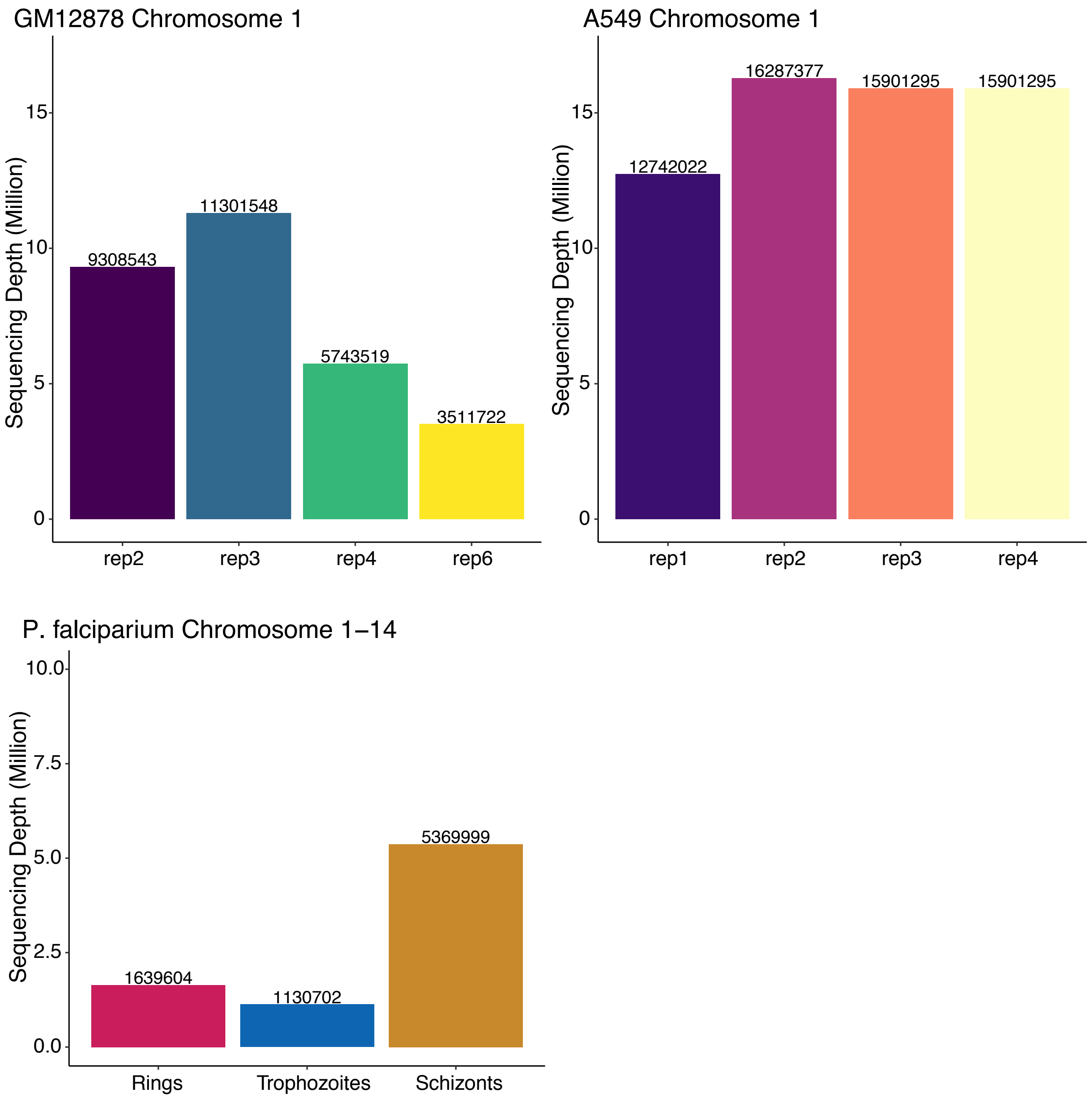


**Supplementary Figure 2 Sequencing depths of the GM12878 and A549 samples on chromosome 1 and *P. falciparum* on chromosome 1-14.** The actual number of valid read-pairs are displayed on the top of each bar.

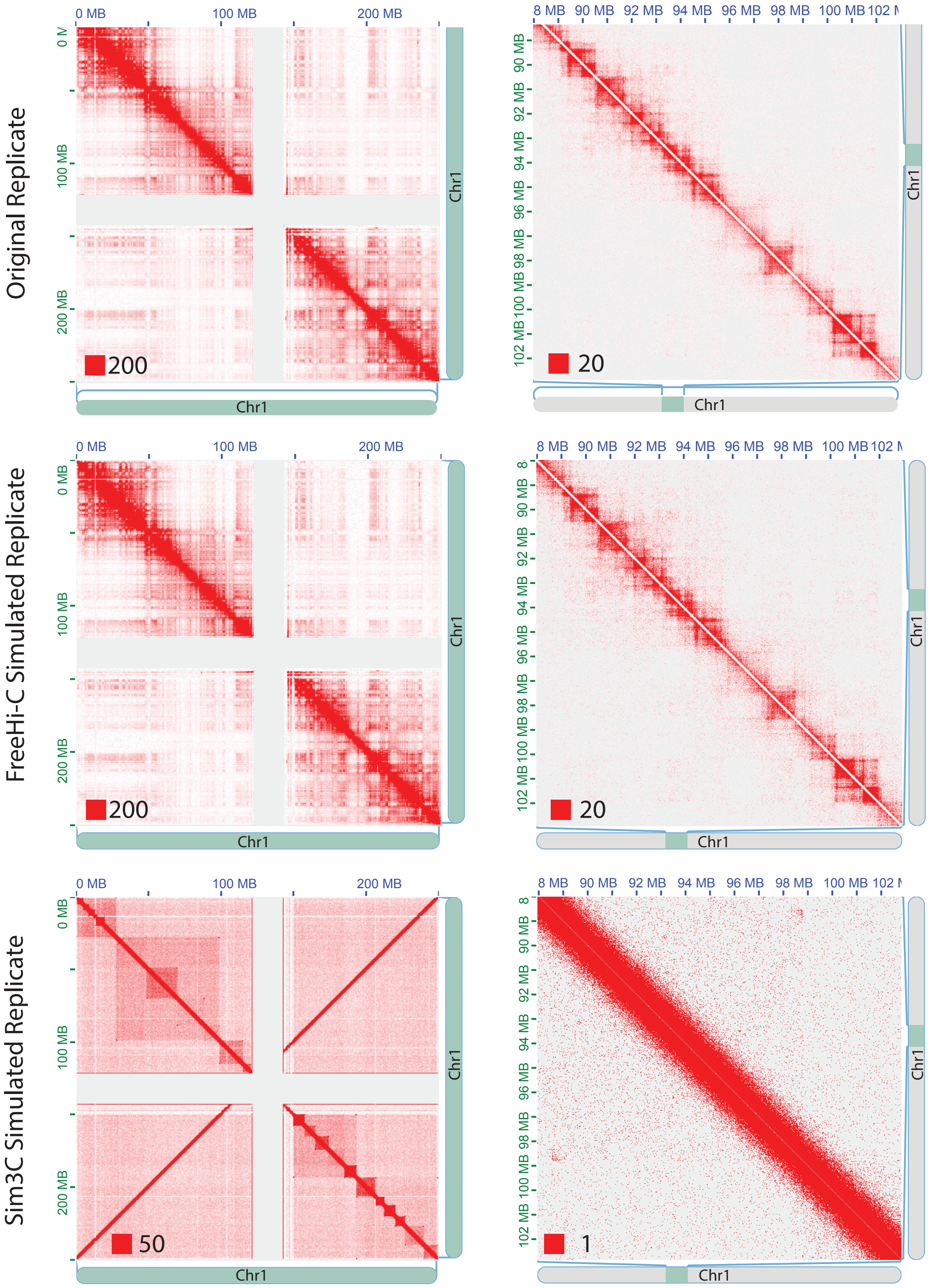
Original Replicate
FreeHi-C Simulated Replicate
Sim3C Simulated Replicate
0 MB
100 MB
200 MB
Chr1
200
20
50
1
90 MB
92 MB
94 MB
96 MB
98 MB
100 MB
102 MB

**Supplementary Figure 3 Hi-C contact matrices of chromosome 1 for rep3 of GM12878.** The first column shows the chromosome-wide Hi-C contact matrices of the original biological replicate and corresponding FreeHi-C and Sim3C simulated replicates with matching sequencing depth of the original replicate. The contact matrices in the second column display a zoom-in at coordinates chr1:88-103Mb. The numbers at the left bottom of each matrix represent the color scale.

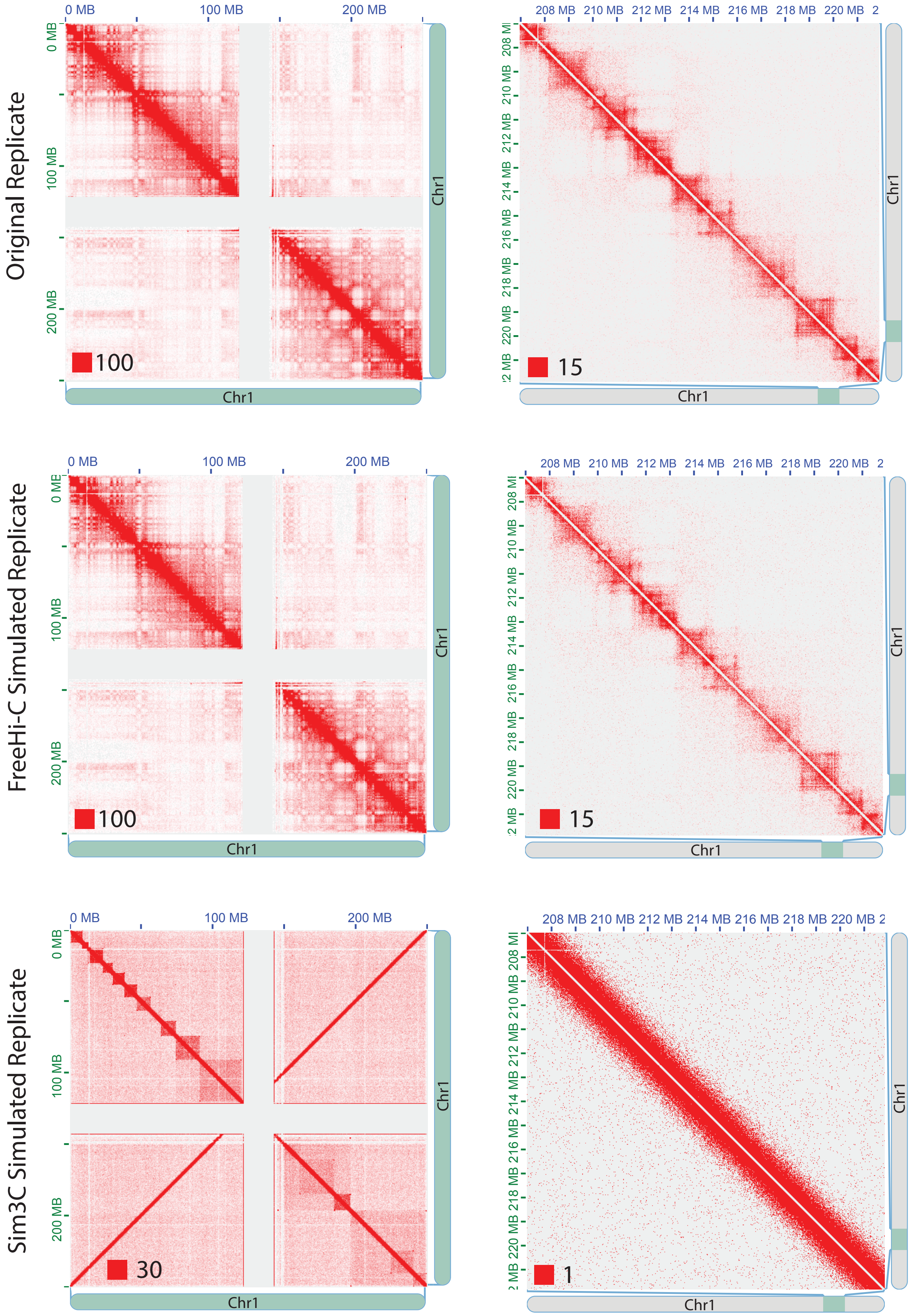
Original Replicate
FreeHi-C Simulated Replicate
Sim3C Simulated Replicate
0 MB
100 MB
200 MB
208 MB
210 MB
212 MB
214 MB
216 MB
218 MB
220 MB
Chr1
100
15
30
1

**Supplementary Figure 4 Hi-C contact matrices of chromosome 1 for rep3 of GM12878.** The first column shows the chromosome-wide Hi-C contact matrices of the original biological replicate and corresponding FreeHi-C and Sim3C simulated replicates with matching sequencing depth of the original replicate. The contact matrices in the second column display a zoom-in at coordinates chr1:207-222Mb. The numbers at the left bottom of each matrix represent the color scale.

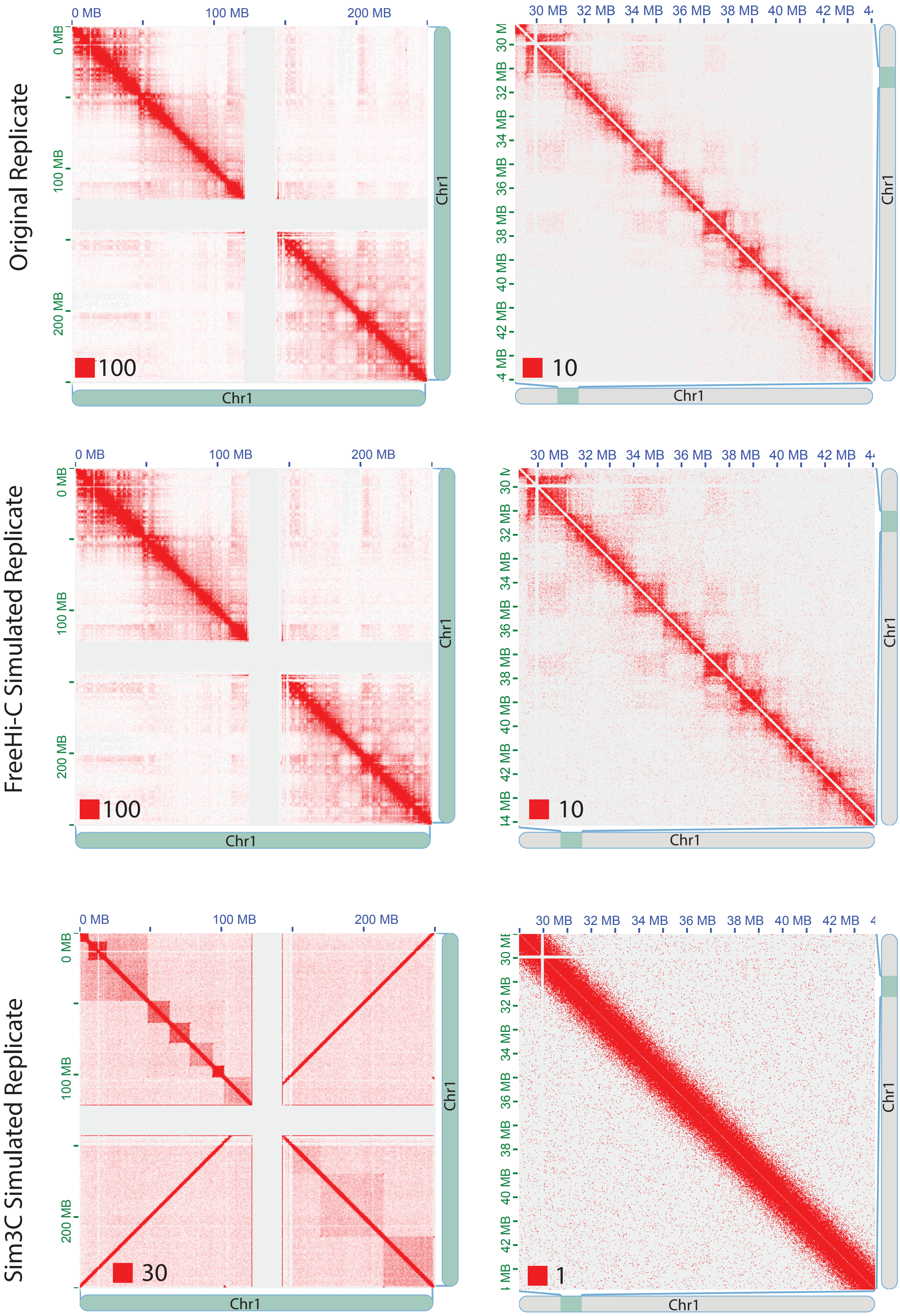
Original Replicate
FreeHi-C Simulated Replicate
Sim3C Simulated Replicate
0 MB
100 MB
200 MB
Chr1
100
10
30
1
30 MB
32 MB
34 MB
36 MB
38 MB
40 MB
42 MB

**Supplementary Figure 5 Hi-C contact matrices of chromosome 1 for rep3 of GM12878.** The first column shows the chromosome-wide Hi-C contact matrices of the original biological replicate and corresponding FreeHi-C and Sim3C simulated replicates with matching sequencing depth of the original replicate. The contact matrices in the second column display a zoom-in at coordinates chr1:29-44Mb. The numbers at the left bottom of each matrix represent the color scale.

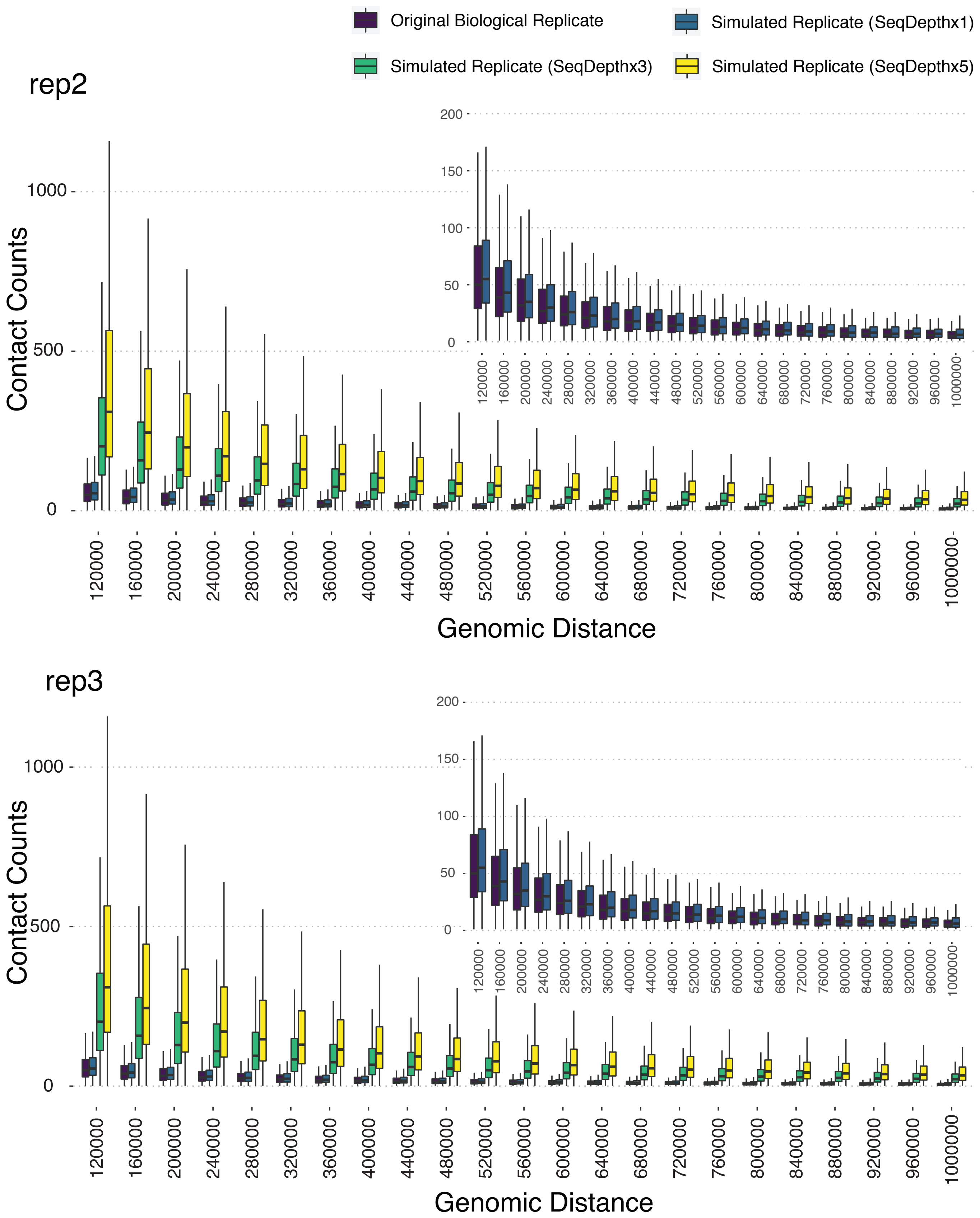


**Supplementary Figure 6 Genomic distance stratified comparison of the Hi-C signal, as quantified by the number of valid read pairs, between the original biological replicates and corresponding FreeHi-C simulated replicates of varying depths for rep2 and rep3 of GM12878.**

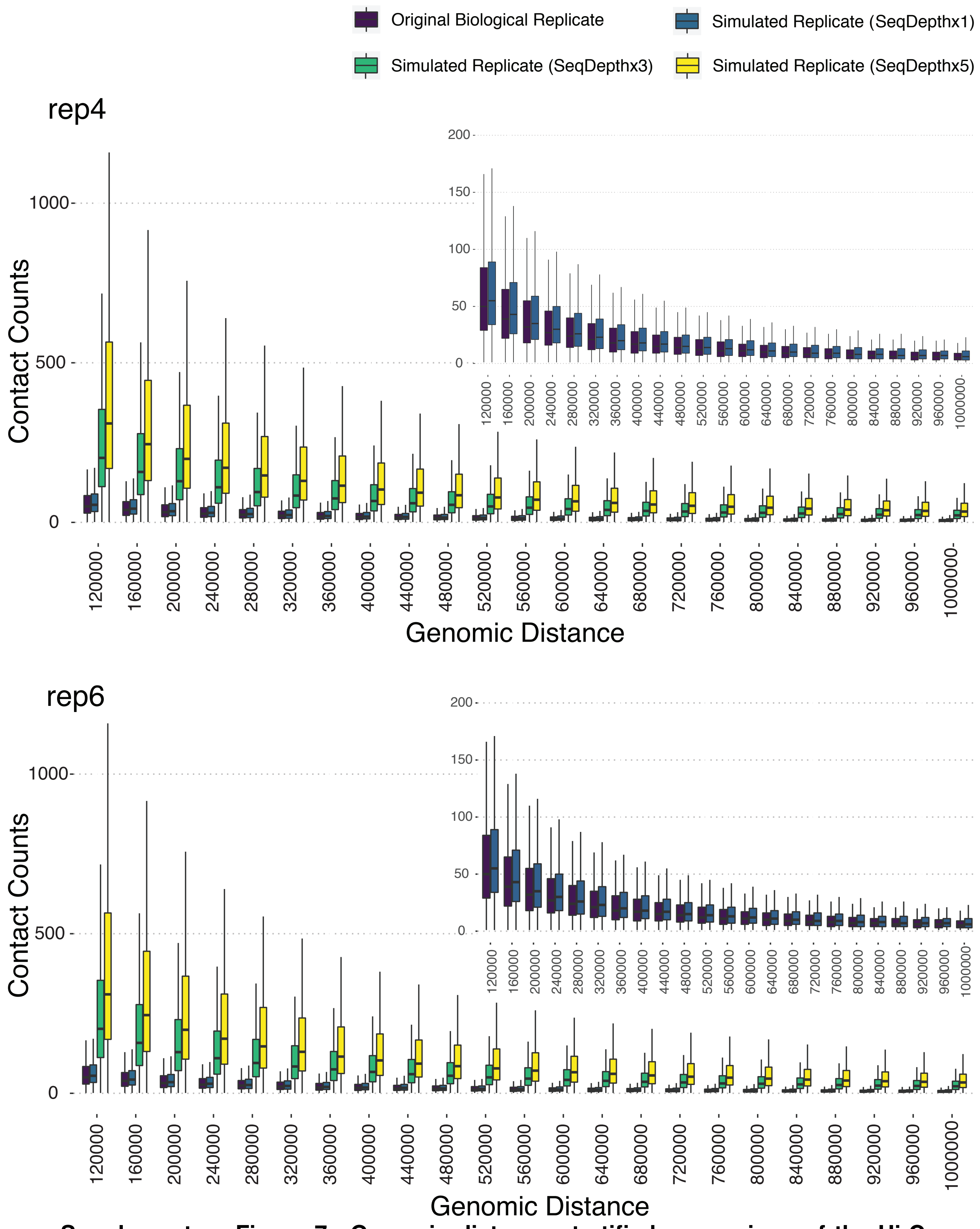


**Supplementary Figure 7 Genomic distance stratified comparison of the Hi-C signal, as quantified by the number of valid read pairs, between the original biological replicates and corresponding FreeHi-C simulated replicates of varying depths for rep4 and rep6 of GM12878.**

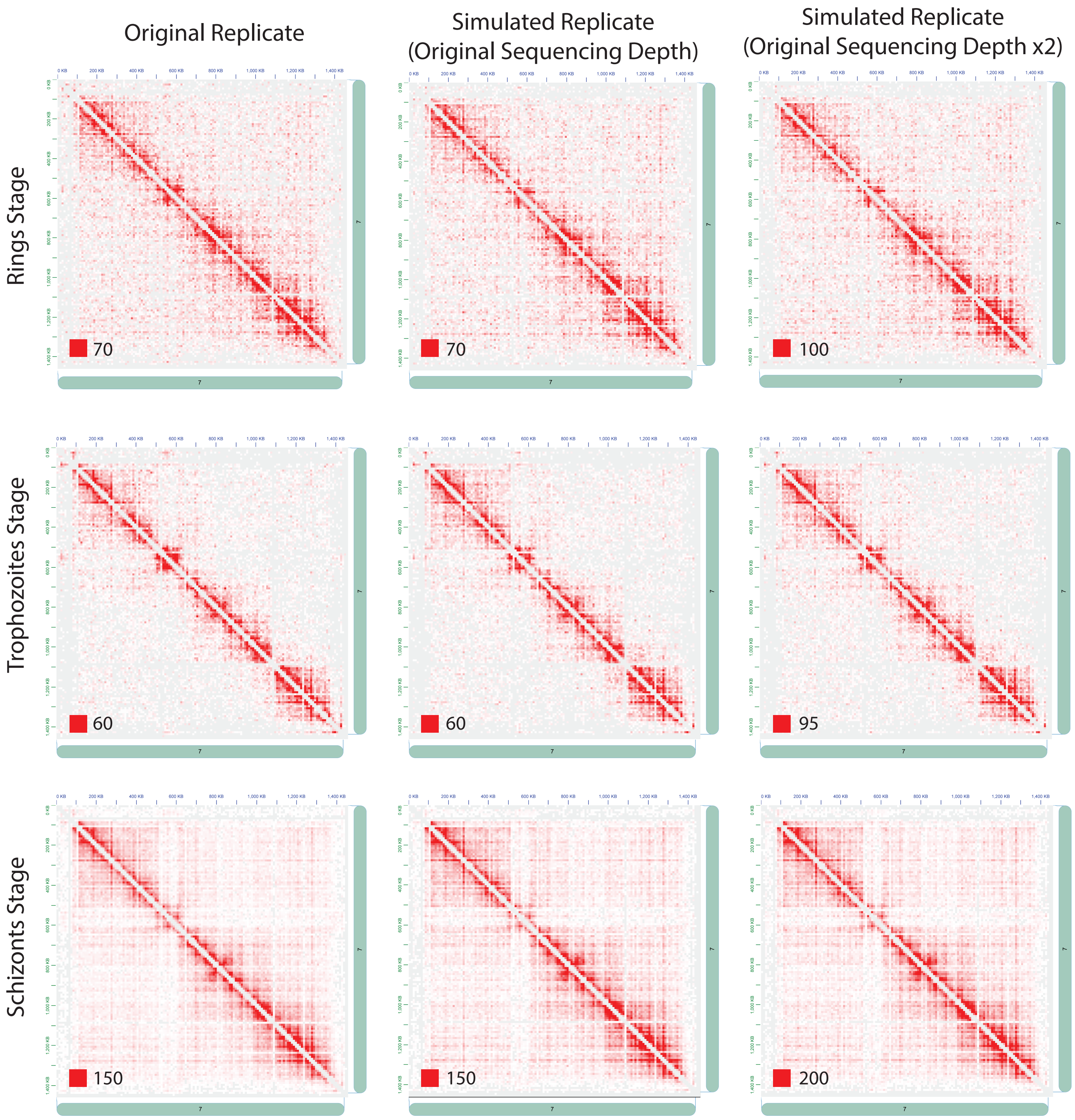


**Supplementary Figure 8 Hi-C contact matrices for chromosome 7 of *P. falciparum* for the Rings, Trophozoites, and Schizonts stages.** Column one corresponds to the original sample and columns two to three are FreeHi-C simulations at the original and twice the depth of the biological sample, respectively. The numbers at the left bottom of each matrix represent the color scale.

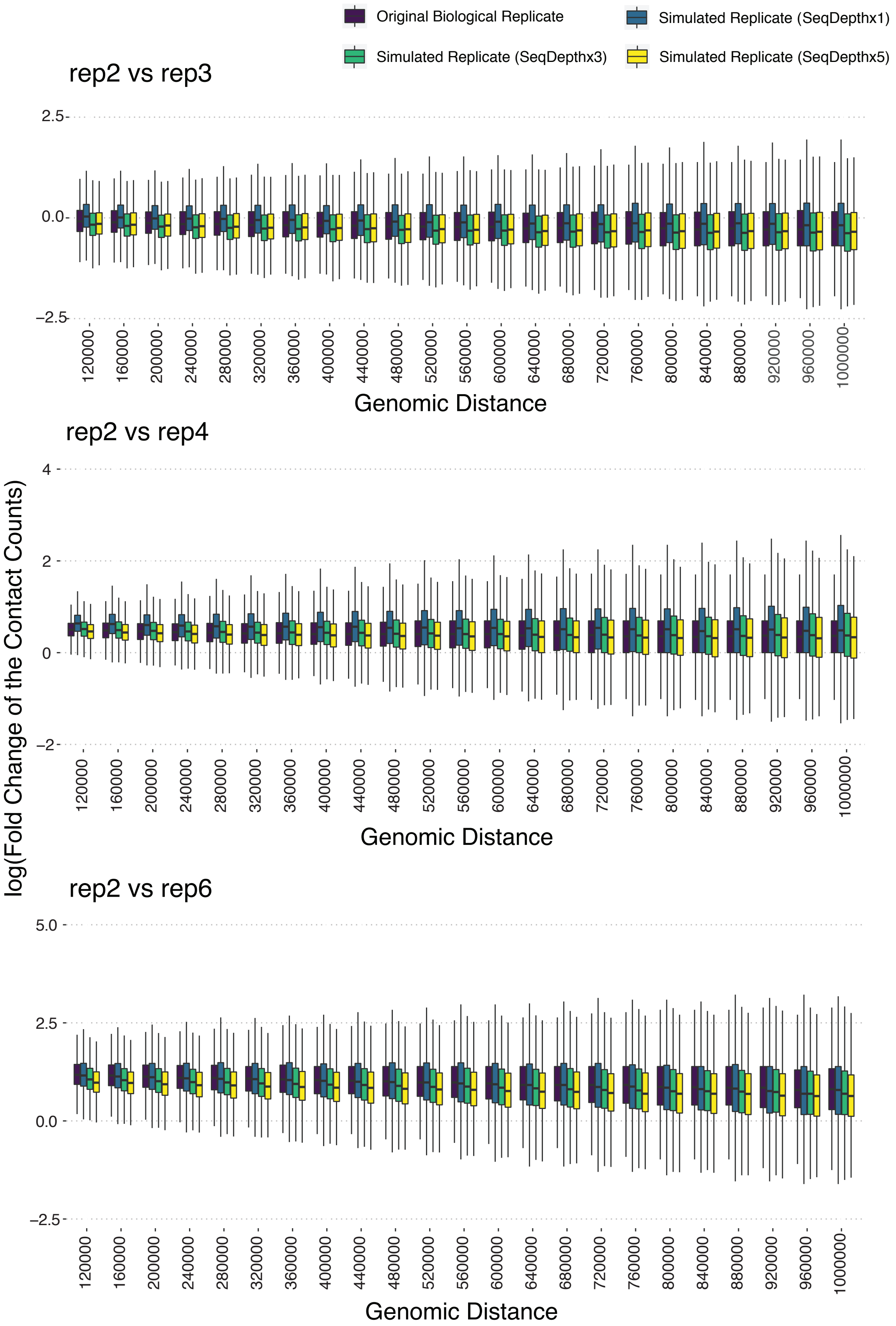


**Supplementary Figure 9 Genomic distance stratified log-fold-change comparison of the Hi-C signal, as quantified by the contact counts, between every pair**

**of GM12878 replicates and the corresponding pair of FreeHi-C simulated replicates.**

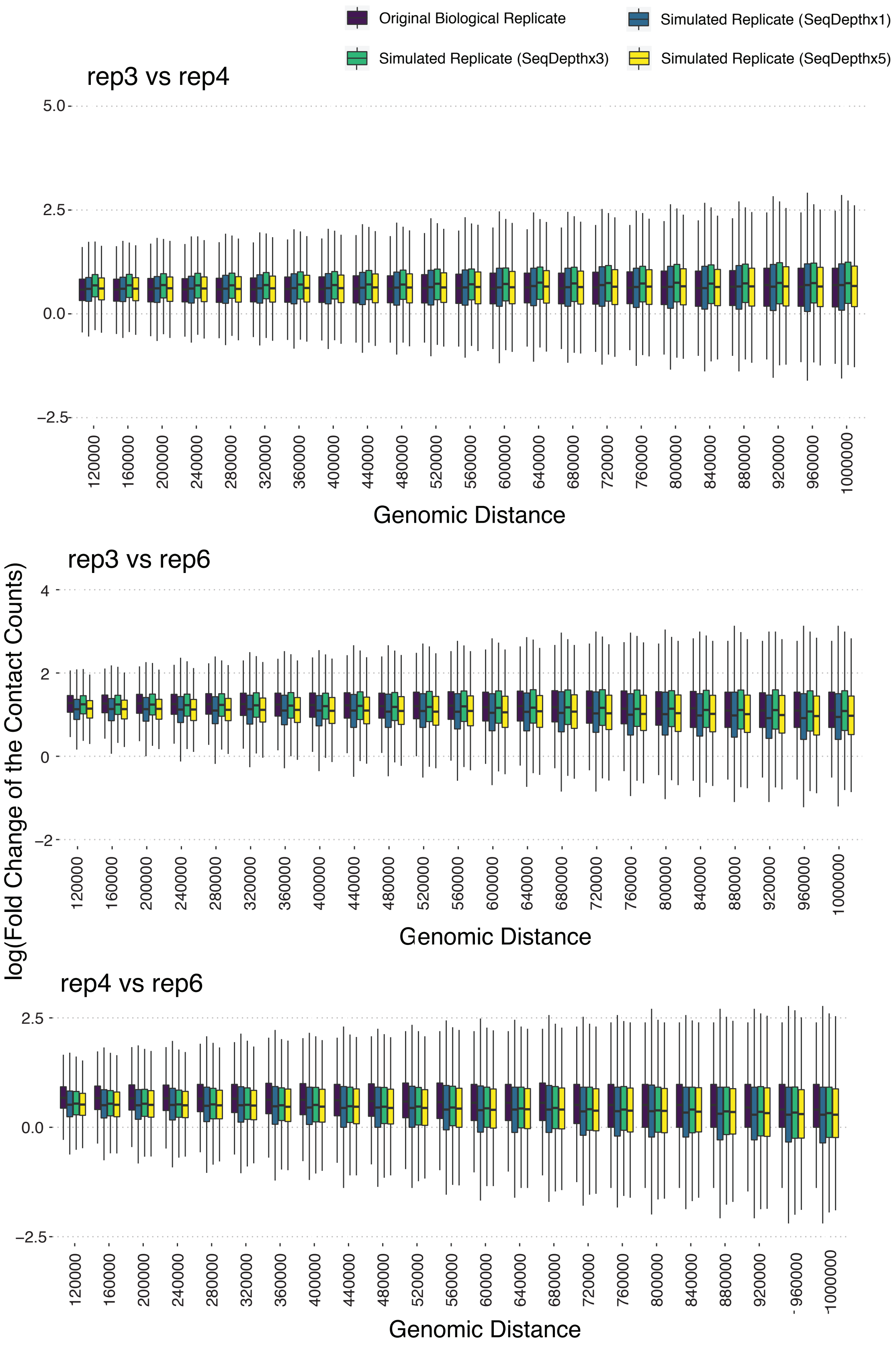


**Supplementary Figure 10 (Cont'd) Genomic distance stratified log-fold-change comparison of the Hi-C signal, as quantified by the contact counts, between ev-**

**ery pair of GM12878 replicates and the corresponding pair of FreeHi-C simulated replicates.**

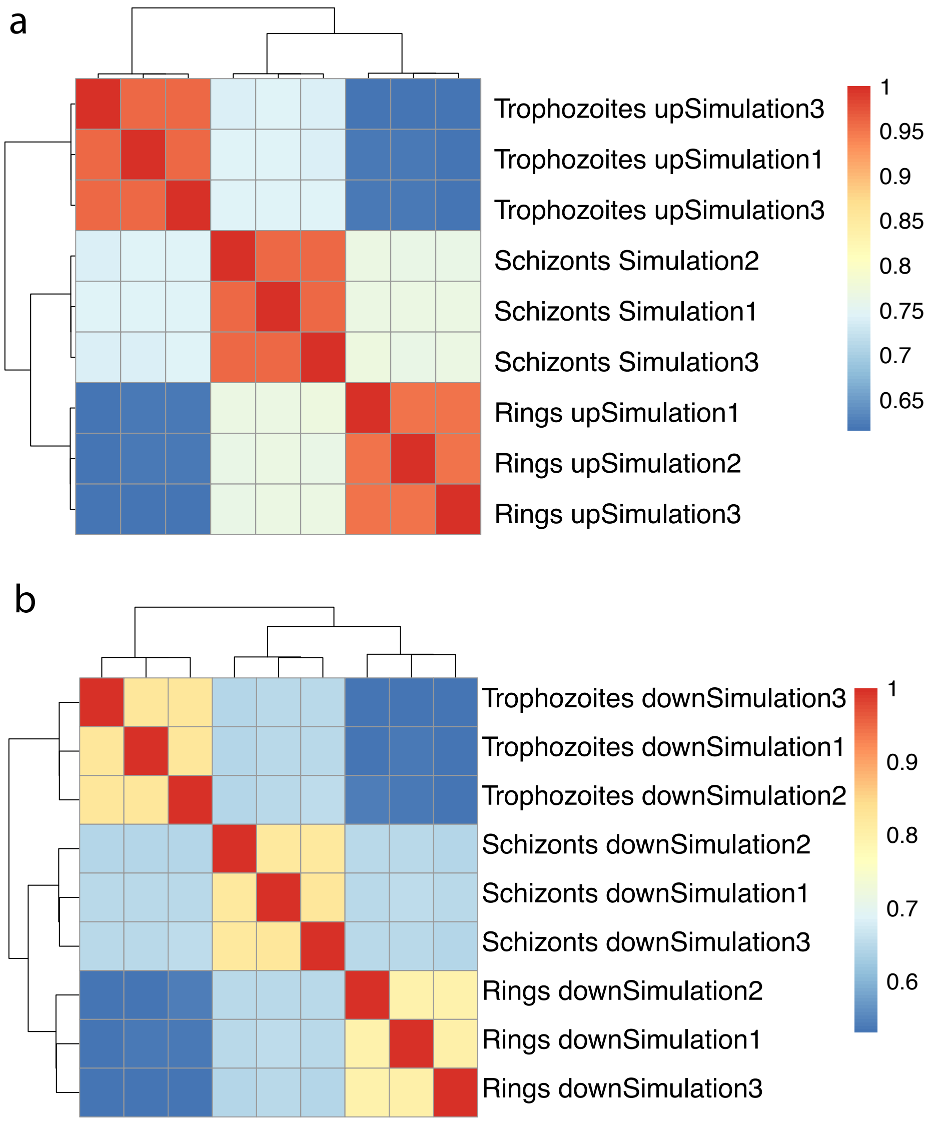


**Supplementary Figure 11 Hierarchical clustering of the FreeHi-C replicates simulated to the same sequencing depth as the original *P. falciparum* Schizonts (a, up-simulation) or Trophozoites (b, down-simulation) stage samples.** Regardless of up (a) or down (b) simulation, FreeHi-C replicates capture the known relationship between the three stages.

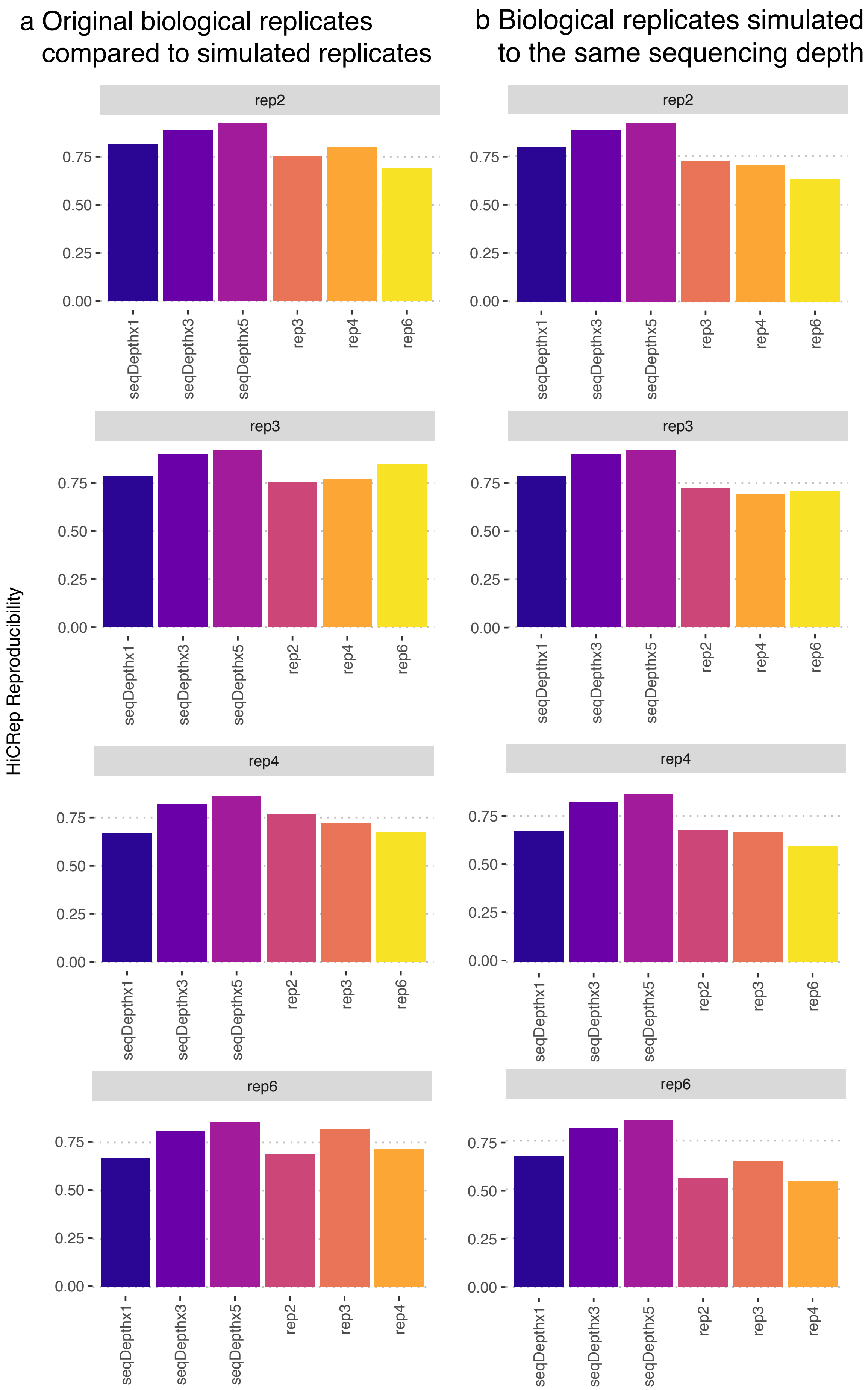
a Original biological replicates compared to simulated replicates
b Biological replicates simulated to the same sequencing depth
HiCRep Reproducibility
rep2
rep3
rep4
rep6
0.00
0.25
0.50
0.75
seqDepthx1
seqDepthx3
seqDepthx5
rep2
rep3
rep4
rep6

**Supplementary Figure 12 HiCRep quantified reproducibility of contact matrices among simulated and biological replicates of GM12878. a.** Reproducibility of the contact matrices between the targeted biological replicates of GM12878 (listed on top of the panel) and corresponding FreeHi-C simulated replicates of varying depths as well as the rest of the biological replicates of GM12878. **b.** Reproducibility of the contact matrices between the targeted biological replicates of GM12878 (listed on the top of the panel) and corresponding FreeHi-C simulated replicates of varying depths as well as FreeHi-C simulated replicates of the rest of the biological samples that are simulated to the same sequencing depth of the targeted biological replicate.

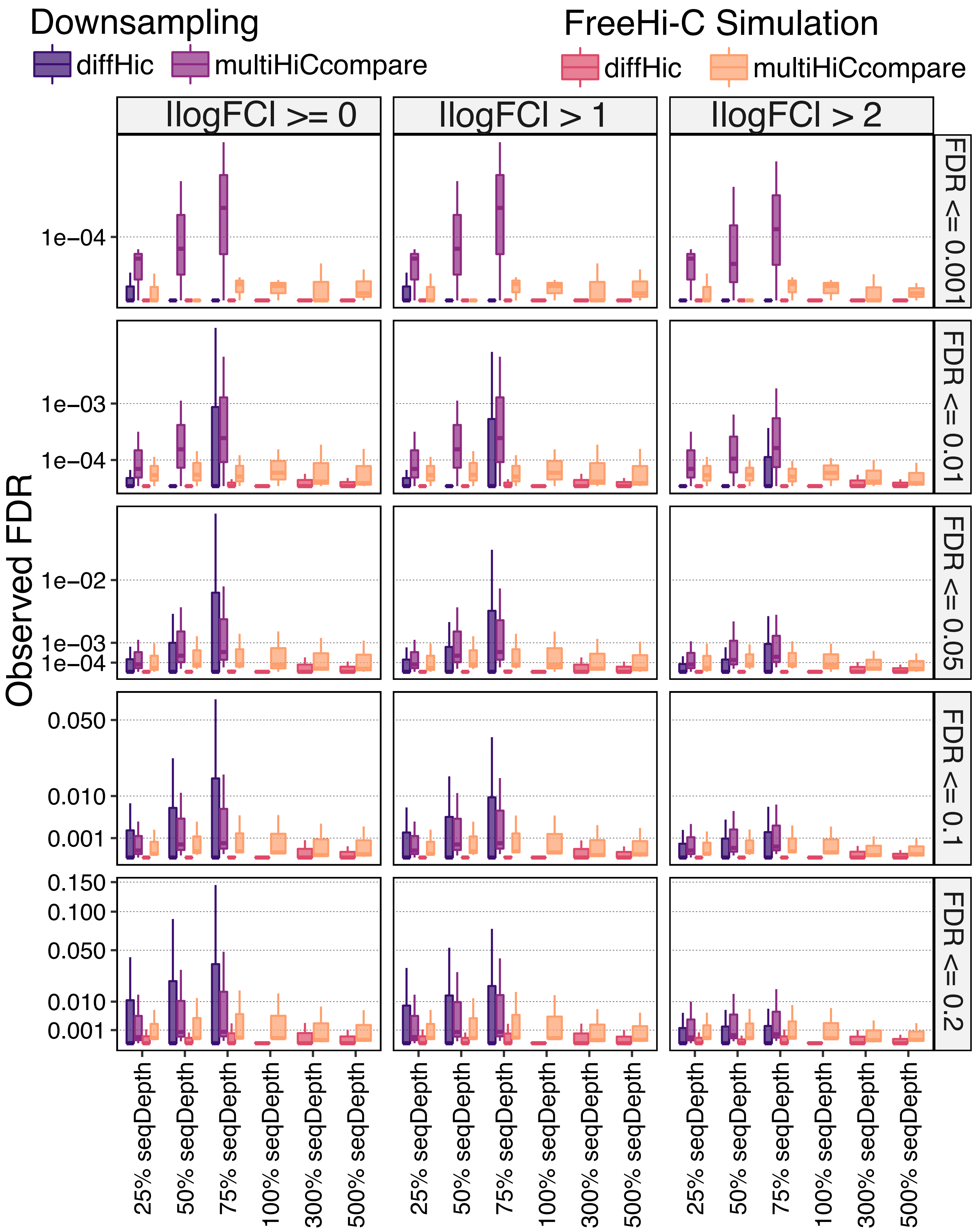

Downsampling
diffHic
multiHiCcompare
FreeHi-C Simulation
diffHic
multiHiCcompare
|logFC| >= 0
|logFC| > 1
|logFC| > 2
FDR <= 0.001
FDR <= 0.01
FDR <= 0.05
FDR <= 0.1
FDR <= 0.2
Observed FDR
1e−04
1e−03
1e−02
0.050
0.010
0.001
0.150
0.100
25% seqDepth
50% seqDepth
75% seqDepth
100% seqDepth
300% seqDepth
500% seqDepth

**Supplementary Figure 13 Differential chromatin interaction detection between biological replicates of GM12878 for evaluating the false discovery rates of diffHic and multiHiCcompare, across a series of sequencing depths using FreeHi-C and downsampling.** Target FDR levels are delineated as the panel labels on the right-hand side. In the downsampling experiments, rep2, rep3, rep4, and rep6 of GM12878 are downsampled to 1/4, 1/2, and 3/4 of the original sequencing depths. Observed false discovery rates of diffHic and multiHiCcompare are depicted in dark purple and light purple, respectively. Biological replicates are also simulated to 1/4, 1/2, 3/4, the same as, three times, and five times the original sequencing depths with FreeHi-C. Resulting false discovery rates of simulated replicates are depicted in red and orange.

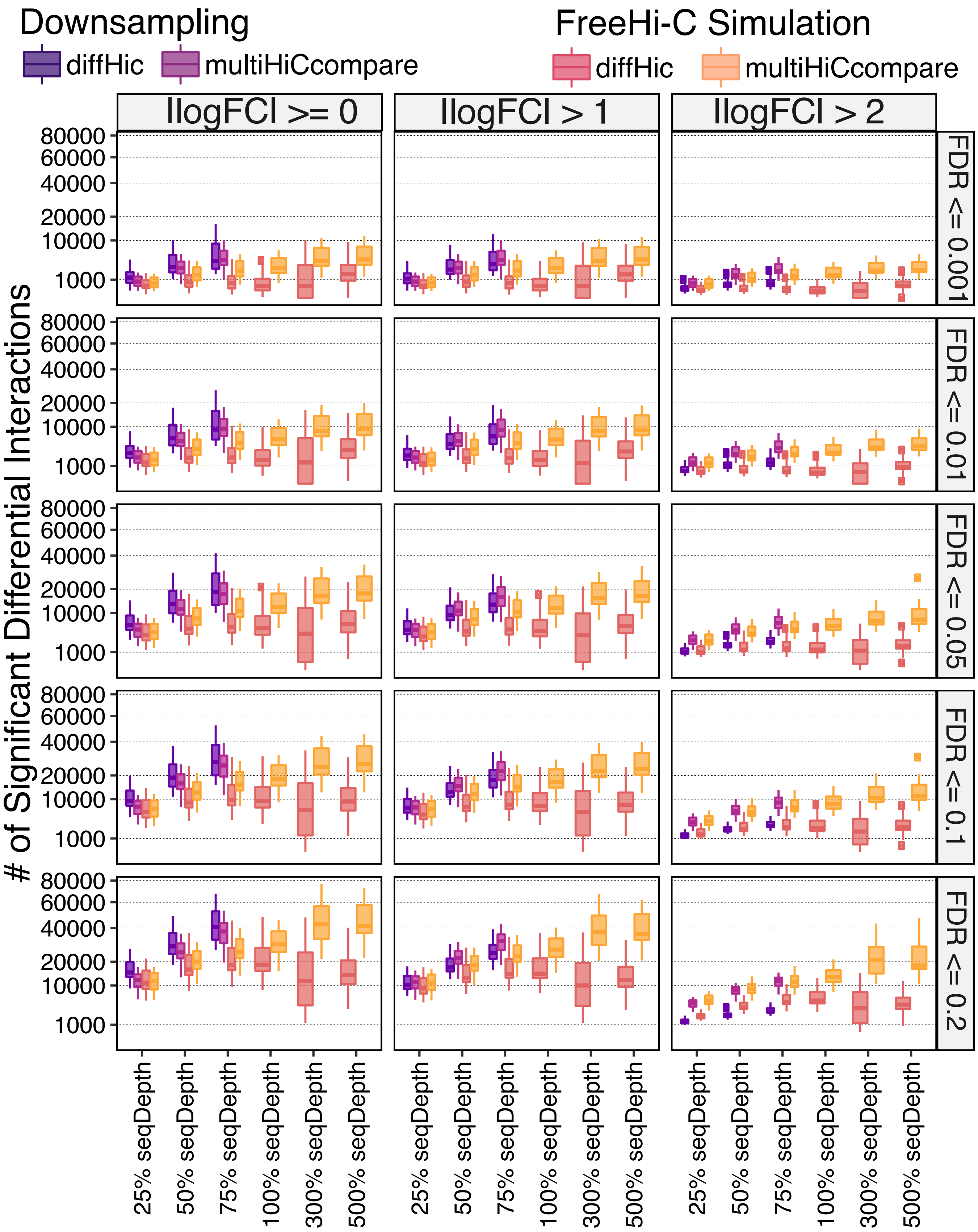
Downsampling
diffHic
multiHiCcompare
FreeHi-C Simulation
diffHic
multiHiCcompare
|logFC| >= 0
|logFC| > 1
|logFC| > 2
FDR <= 0.001
FDR <= 0.01
FDR <= 0.05
FDR <= 0.1
FDR <= 0.2
### of Significant Differential Interactions
80000
60000
40000
20000
10000
1000
25% seqDepth
50% seqDepth
75% seqDepth
100% seqDepth
300% seqDepth
500% seqDepth

**Supplementary Figure 14 Differential chromatin interaction detection between two replicates of GM12878 and two replicates of A549 for evaluating the power of diffHic and multiHiCcompare, across a series of sequencing depths with FreeHi-C and downsampling.** Target FDR levels for DCI are delineated as the panel labels on the right-hand side. In the downsampling experiments, rep2, rep3, rep4, and rep6 of GM12878 and rep1-rep4 of A549 are downsampled to 1/4, 1/2, and 3/4 of the original sequencing depths. The numbers of significant DCIs detected for diffHic, and multiHiCcompare are depicted in dark purple and light purple, respectively. Biological replicates of both cell lines are also simulated to 1/4, 1/2, 3/4, the same as, three times, and five times the original sequencing depths with FreeHi-C. Resulting numbers of detected DCIs are depicted in red and orange.

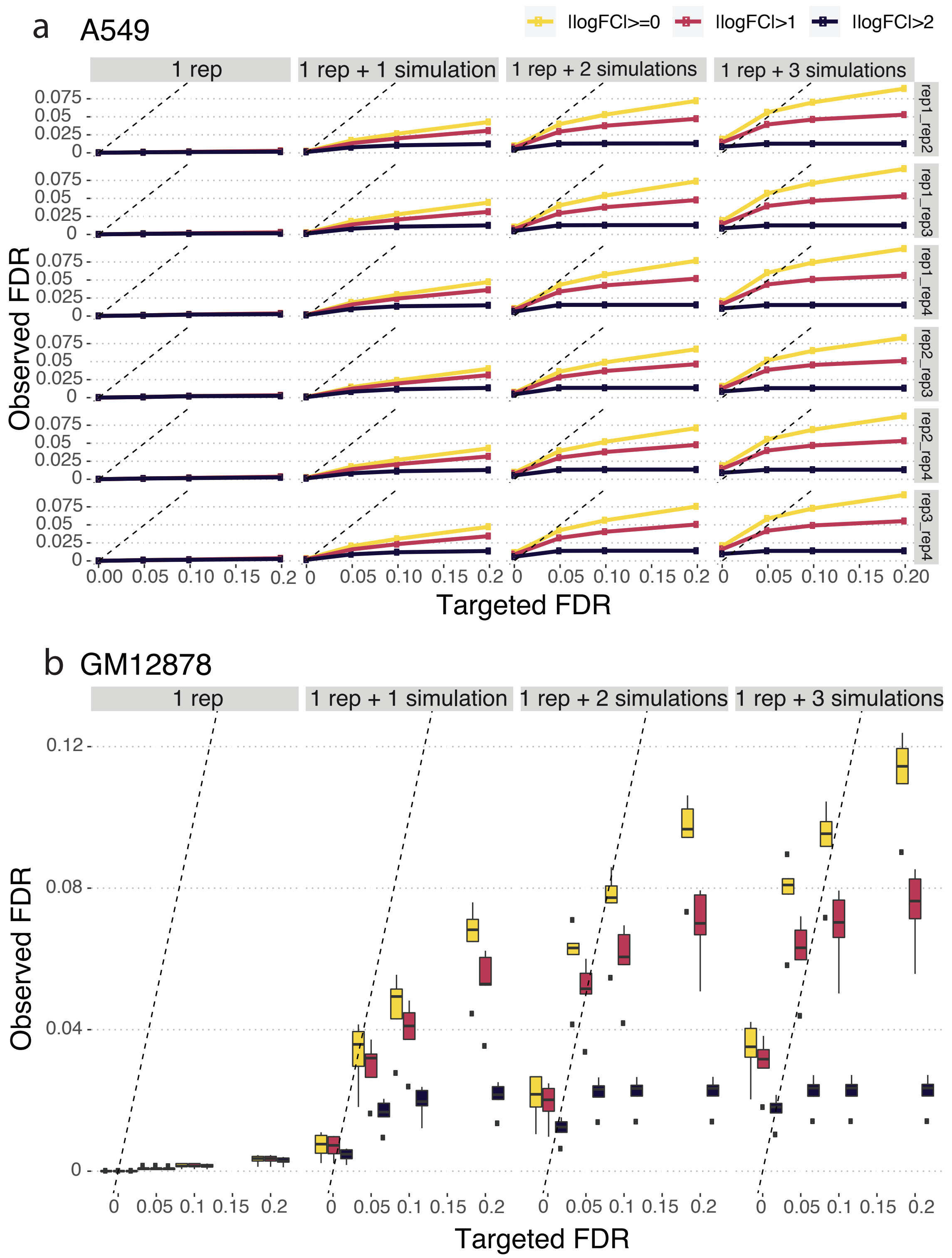


**Supplementary Figure 15 False discovery rate control for differential chromatin interaction detection within pairs of A549 or GM12878 replicates with or without FreeHi-C augmentation. a** Observed versus targeted FDR for differential chromatin interaction detection between pairs of A549 replicates (first column) and augmentation with 1 to 3 FreeHi-C simulated replicates (columns 2-4). These data depict the individual pairs of replicates summarized in Figure 2a. **b** False discovery rate control as in Figure 2a with the GM12878 replicates. Dashed lines are y = x.

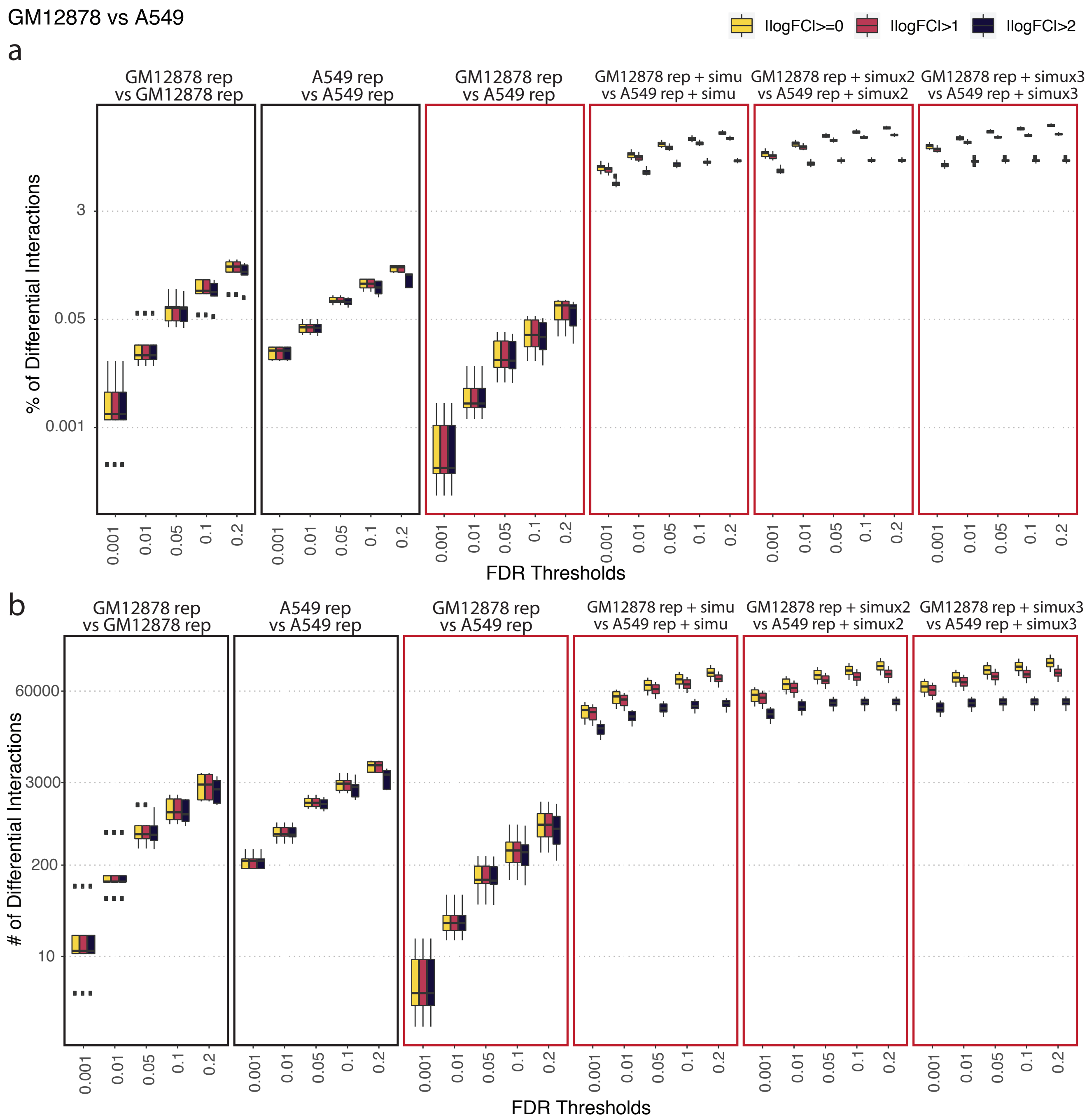


**Supplementary Figure 16 Power of differential chromatin interaction detection between pairs of A549 or GM12878 replicates with or without FreeHi-C data augmentation across a sequence of FDR thresholds.** The first two columns are provided as a reference to highlight that with one replicate per condition, the number of differential chromatin interactions detected between conditions (third column) can be smaller than that of within conditions (number of false discoveries in the first two columns). Columns 4 to 6 illustrate the striking increase in the numbers of detected differential chromatin interactions with FreeHi-C data augmentation. The y-axes are on a log scale. Panels **a** and **b** report in the y-axes the percentage of candidate differential chromatin interactions and number of differential chromatin interactions, respectively.

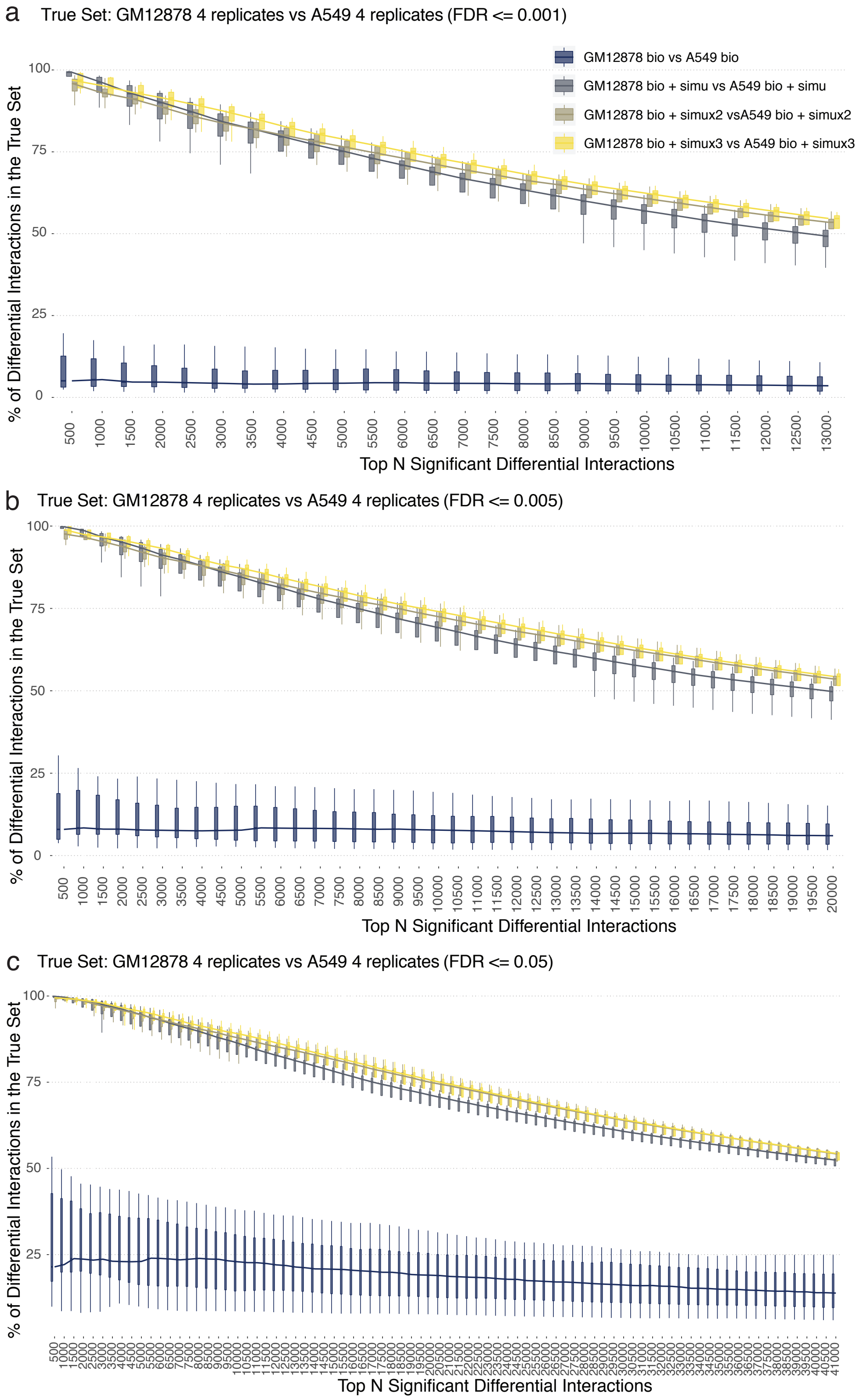


**Supplementary Figure 17 Percentage of differential chromatin interactions (DCIs), detected under the one replicate per condition of GM12878 versus A549 comparison with and without FreeHi-C data augmentation, among the gold standard DCIs set.** True set of DCIs are identified at FDR of 0.001 (**a**), 0.005 (**b**) and 0.05 (**c**).

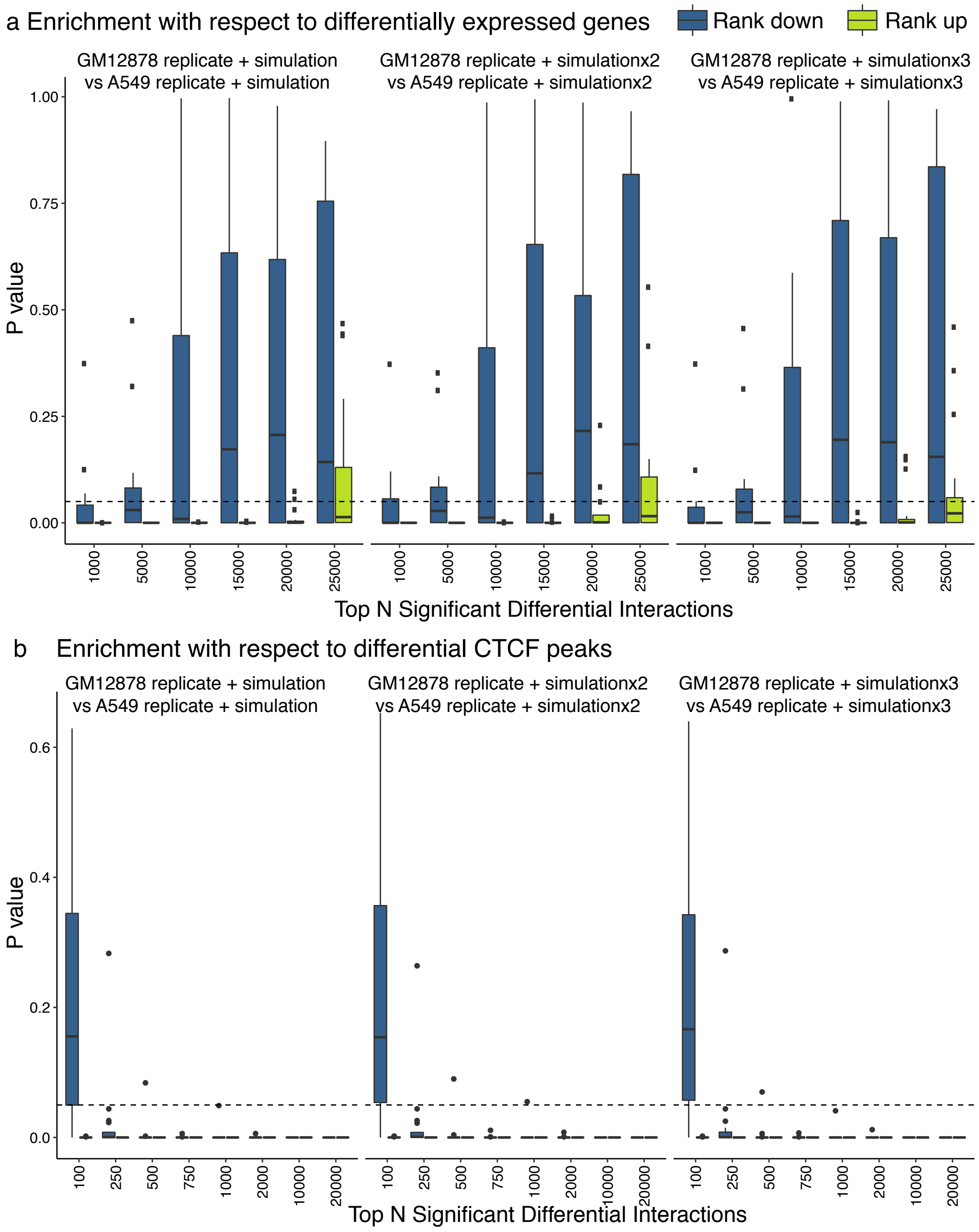


**Supplementary Figure 18 Evaluation of detected differential chromatin interactions by external RNA-seq and CTCF ChIP-seq data of GM12878 and A549 cells.** **a** Ranked up (green) DCIs of the one biological replicate per condition setting as a result of FreeHi-C data augmentation are significantly enriched for differentially expressed (DE) genes. In contrast, DCIs that are ranked lower (blue) among the top N DCIs, N = 1000, 5000,..., 25000, as a result of FreeHi-C augmentation do not significantly overlap with the DE genes. **b** Ranked up (green) and down (blue) DCIs of the one biological replicate per condition setting as a result of FreeHi-C data augmentation are significantly enriched for differential CTCF peaks. Both sets show significant enrichment for CTCF peaks, with the exception of significant enrichment for the ranked up DCIs compared to non-enrichment of ranked down DCIs for differential CTCF peaks among the top 100 DCIs. Dashed lines depict p-value cutoff of 0.05.

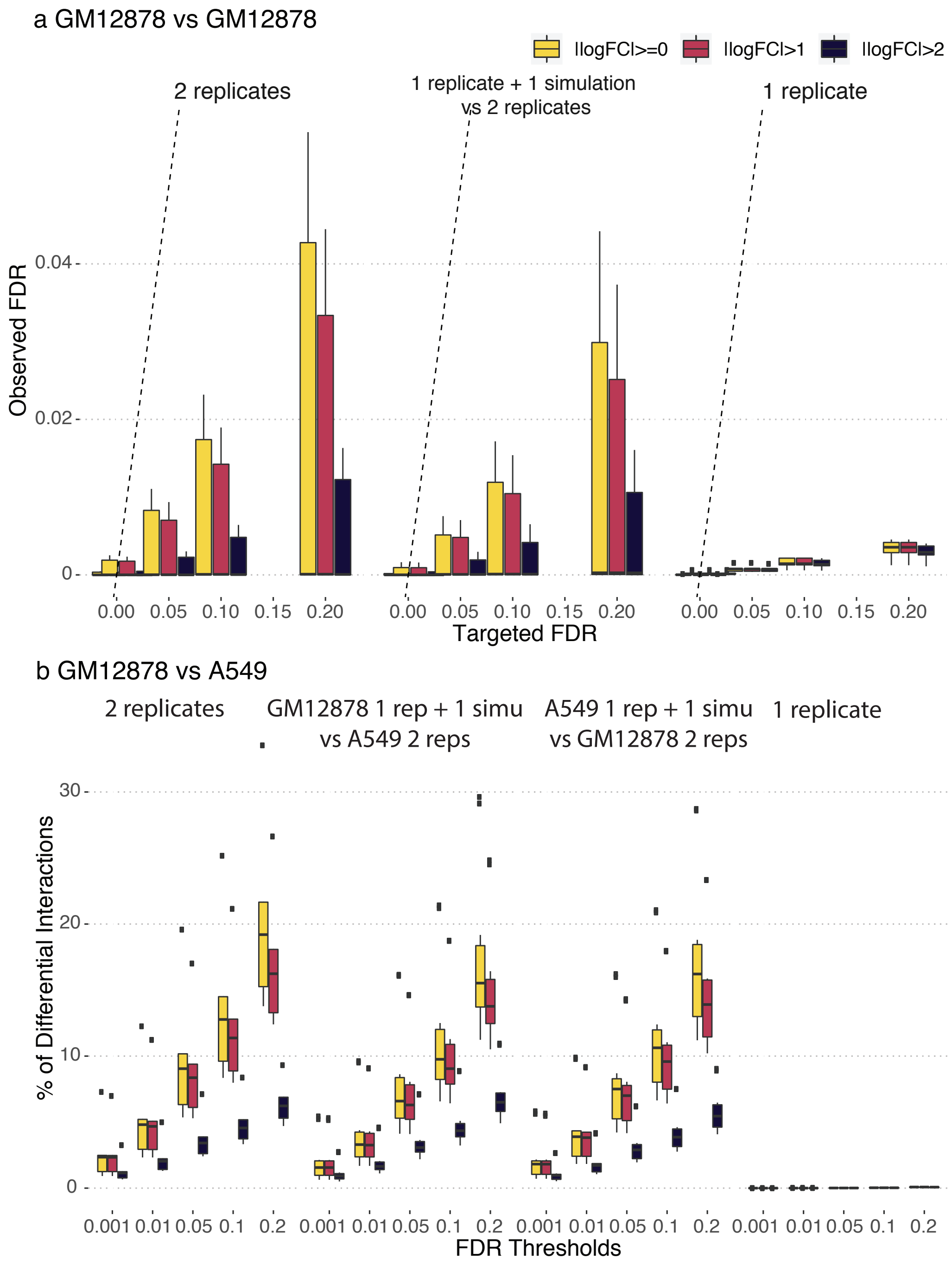


**Supplementary Figure 19 False discovery rate control and power for differential chromatin interaction detection for the uneven numbers of replicates per condition setting. a** Observed versus targeted FDR for differential chromatin interaction detection under variations of replicates per condition (left: two biological replicates per condition; right: one biological replicate per condition; middle: two biological replicates for one condition and FreeHi-C augmentation of the single biological replicate of the other condition). Middle panel corresponds to data augmentation with FreeHi-C and exhibits FDR control. **b** Power, presented as the percentage of candidate DCIs declared as significant at the given FDR threshold, under variations on replicates per condition. Power with two replicates per condition (left panel) depicts the gold standard in this setting.

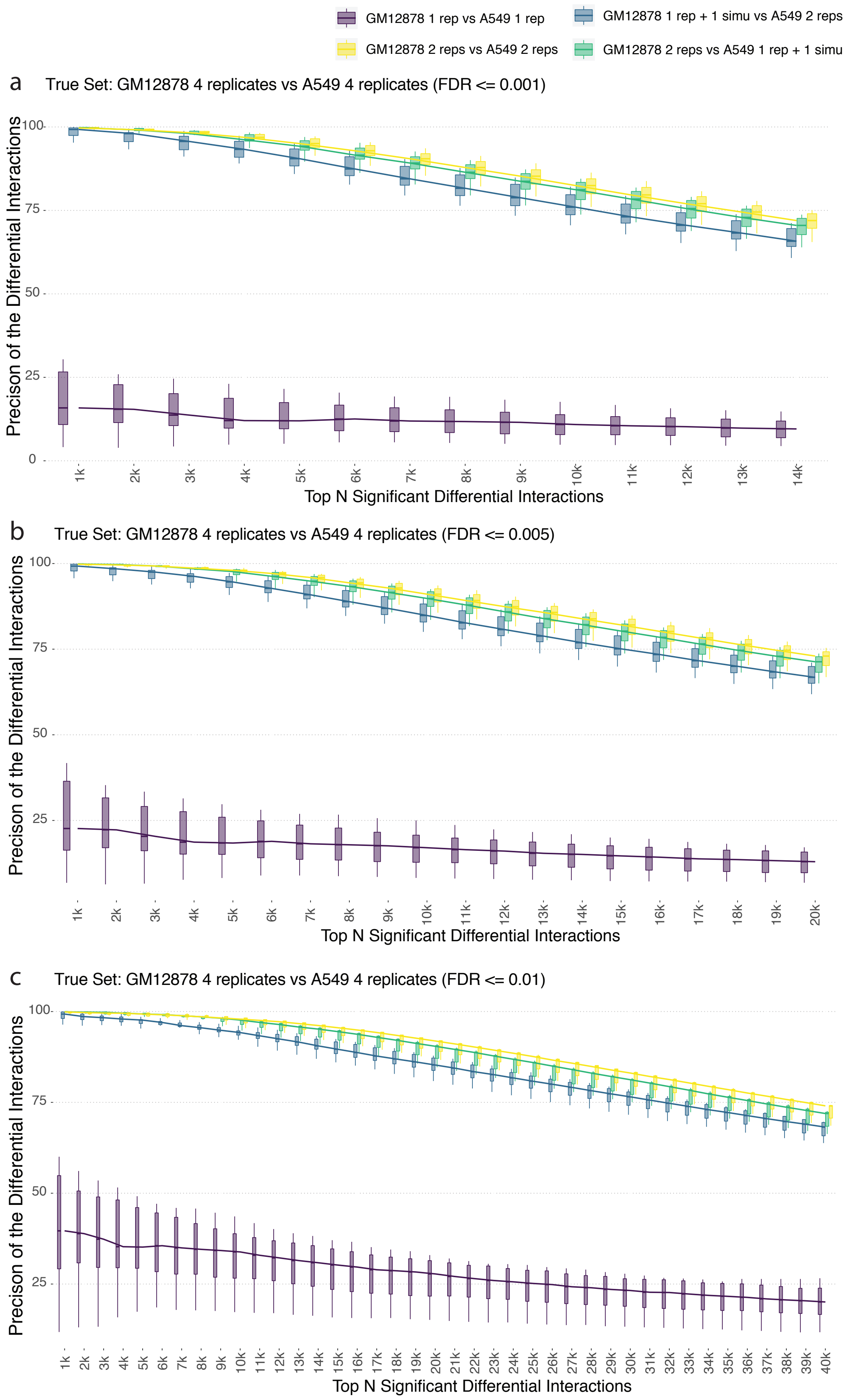

GM12878 1 rep vs A549 1 rep
GM12878 1 rep + 1 simu vs A549 2 reps
GM12878 2 reps vs A549 2 reps
GM12878 2 reps vs A549 1 rep + 1 simu
a
True Set: GM12878 4 replicates vs A549 4 replicates (FDR <= 0.001)
Precison of the Differential Interactions
Top N Significant Differential Interactions
b
True Set: GM12878 4 replicates vs A549 4 replicates (FDR <= 0.005)
Precison of the Differential Interactions
Top N Significant Differential Interactions
c
True Set: GM12878 4 replicates vs A549 4 replicates (FDR <= 0.01)
Precison of the Differential Interactions
Top N Significant Differential Interactions

**Supplementary Figure 20 Percentage of top significant DCIs in the "true" differential chromatin interaction list, that is based on the comparison of the full set of 4 replicates of GM12878 with 4 replicates of A549, under one replicate per condition and the corresponding FreeHi-C data augmentation settings.** The "true" differential chromatin interaction set is defined as the most significant interactions in the comparison of 4 replicates of GM12878 with 4 replicates of A549 thresholded at FDR of 0.001 (**a**), 0.005 (**b**), and 0.05 (**c**). The two biological replicates per condition are included as an upper bound (i.e., the best achievable precision if there was one more biological replicate per condition) for the FreeHi-C data augmentation of single replicates. Leveraging FreeHi-C replicates significantly boosts the precision of top significant DCIs.

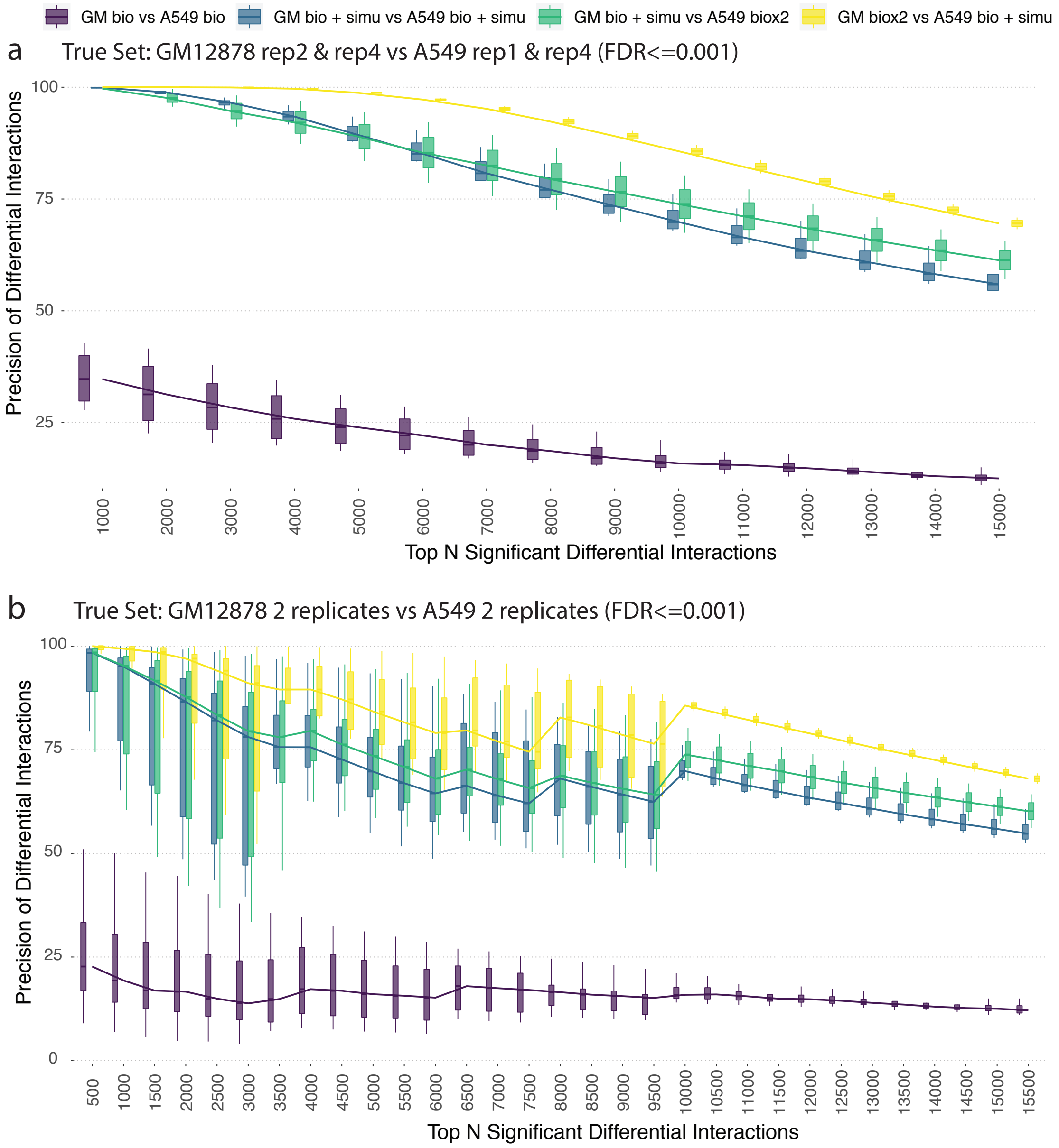


**Supplementary Figure 21 Percentage of top significant DCIs in the "true" differential chromatin interaction list, which is based on the comparison of 2 replicates of GM12878 with 2 replicates of A549, under uneven numbers of replicates condition and the corresponding FreeHi-C data augmentation settings. a** The "true" differential chromatin interaction set is defined as the most significant interactions in the comparison of rep2 and rep4 of GM12878 with rep1 and rep4 of A549 thresholded at FDR 0.001. Differential interaction analysis was carried out considering the following different settings. purple: one replicate out of rep2 and rep4 of GM12878 versus one out of rep1 and rep4 of A549; blue: one replicate out of rep2 and rep4 of GM12878 and its FreeHi-C augmentation versus one out of rep1 and rep4 of A549 and its FreeHi-C augmentation ; green: one replicate out of rep2 and rep4 of GM12878 and its FreeHi-C augmentation versus rep1 and rep4 of A459; yellow: rep2 and rep4 of GM12878 versus one replicate out of rep1 and rep4 of A459 and its FreeHi-C augmentation. **b.** The "true" differential chromatin interaction set is defined as the most significant interactions in the comparison of two replicates of GM12878 with two replicates of A549 thresholded at FDR of 0.001. This is an aggregated version of the precision rates in **a** using all combinations of true sets.

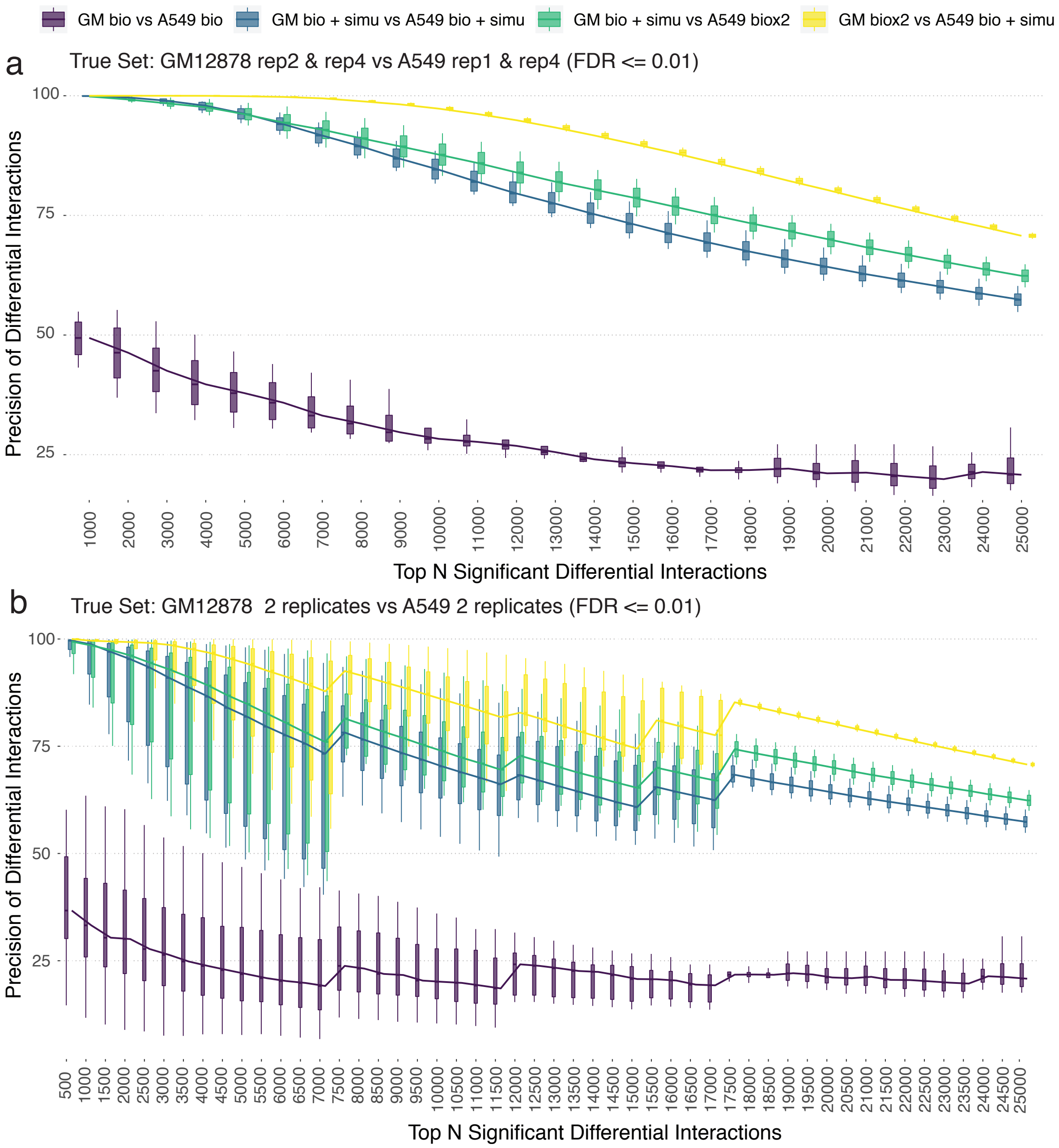


**Supplementary Figure 22** **Same as Supplementary Fig. 21 but with a different FDR threshold of 0.01 for obtaining the gold standard set.**

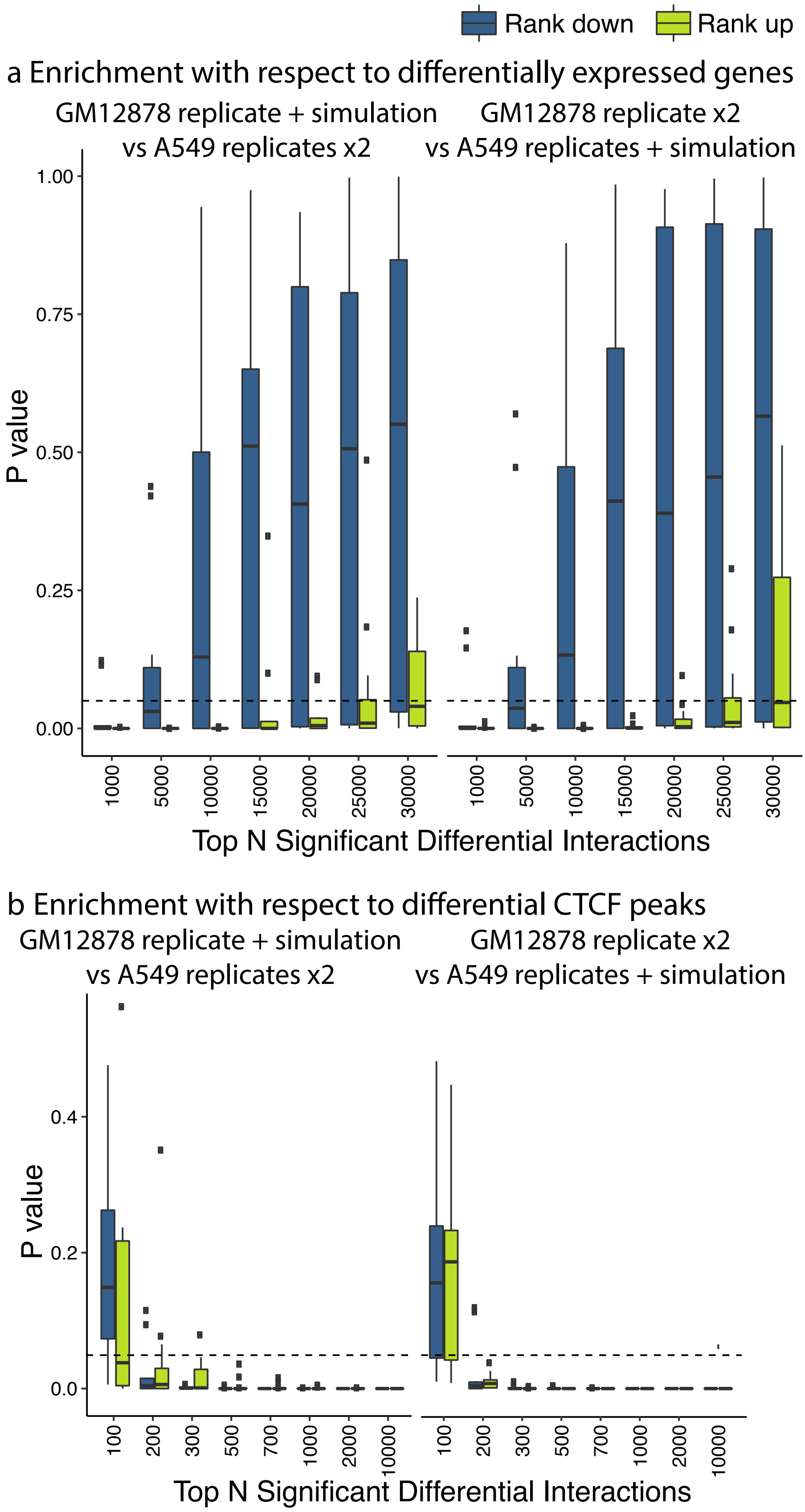


**Supplementary Figure 23 Evaluation of detected differential chromatin interactions by external RNA-seq and CTCF ChIP-seq data of GM12878 and A549 cells.**

**a** Ranked up (green) DCIs of the uneven number of replicates per condition setting as a result of FreeHi-C data augmentation are significantly enriched for differentially expressed (DE) genes. In contrast, DCIs that are ranked lower (blue) among the top N DCIs, N = 1000, 5000,..., 30000, as a result of FreeHi-C augmentation do not significantly overlap with the DE genes. **b** Ranked up (green) and down (blue) DCIs of the one biological replicate per condition setting as a result of FreeHi-C data augmentation are significantly enriched for differential CTCF peaks. Both sets show significant enrichment for CTCF peaks, with the exception at the top 100 DCIs. Dashed lines depict p-value cutoff of 0.05.

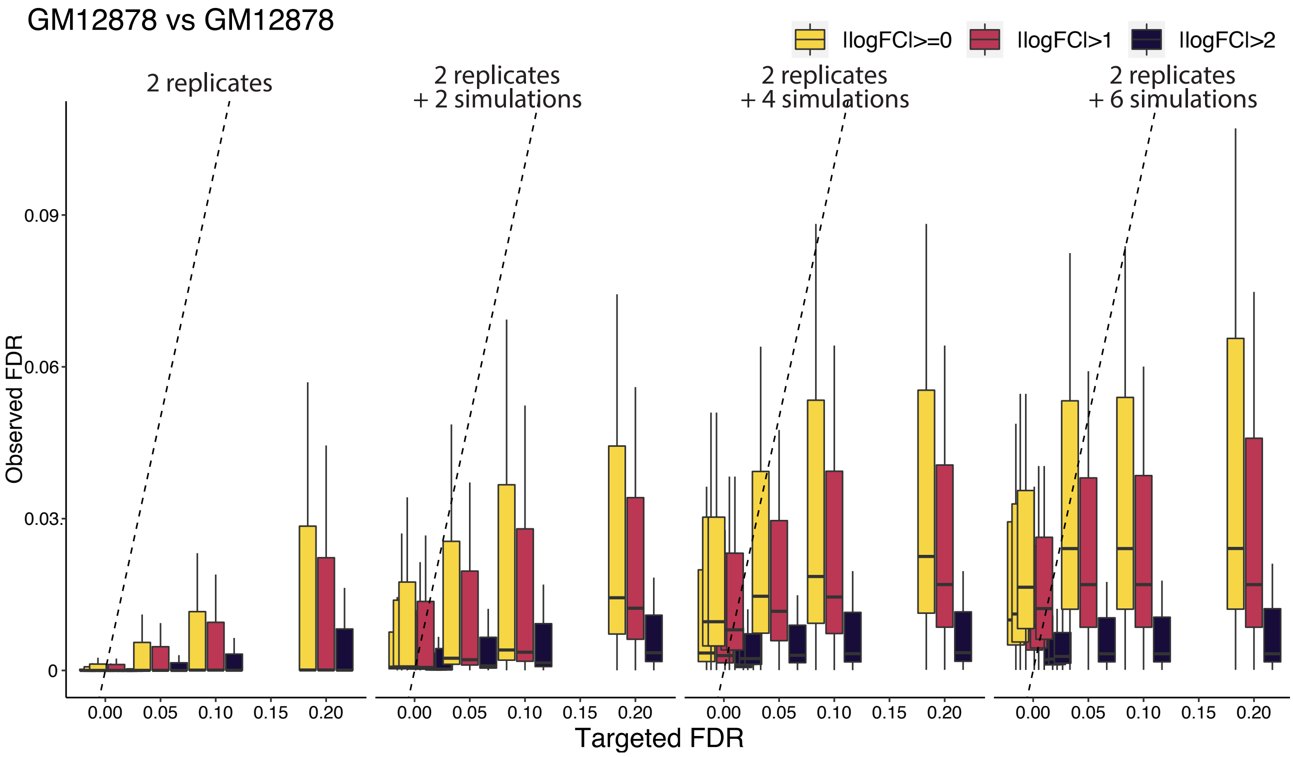


**Supplementary Figure 24 Data augmentation with FreeHi-C replicates preserve false discovery rate control.** Observed false discovery rates of within sample comparisons for GM12878 data (i.e., comparisons of two biological replicates of GM12878 with another set of two biological replicates from GM12878). This is an analog of Figure 2e for GM12878.

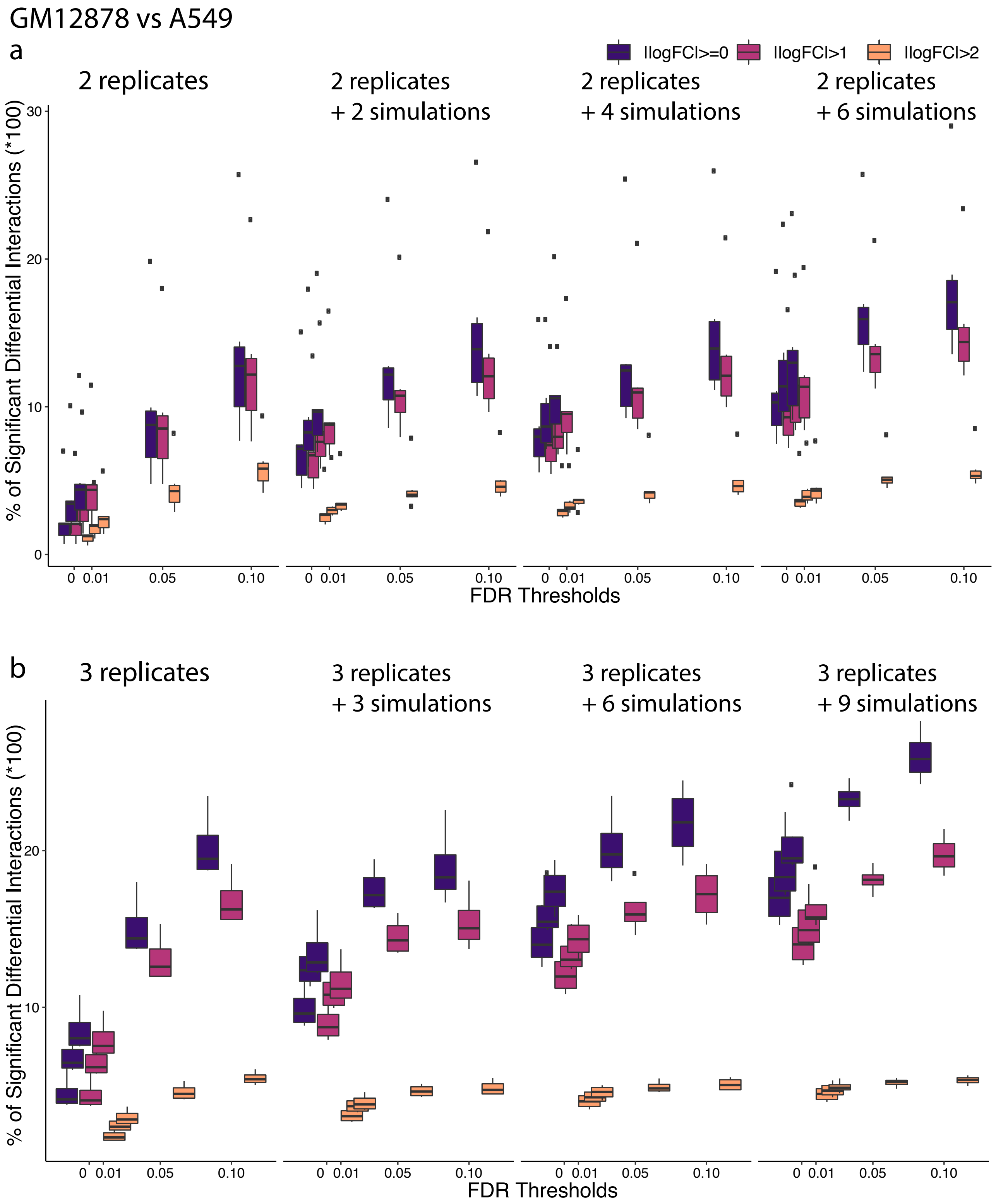


**Supplementary Figure 25 Power for differential chromatin interaction detection between GM12878 and A459 samples under a variety of settings with and without FreeHi-C data augmentation. a** and **b** delineate cases with 2 and 3 biological replicates per condition, respectively.

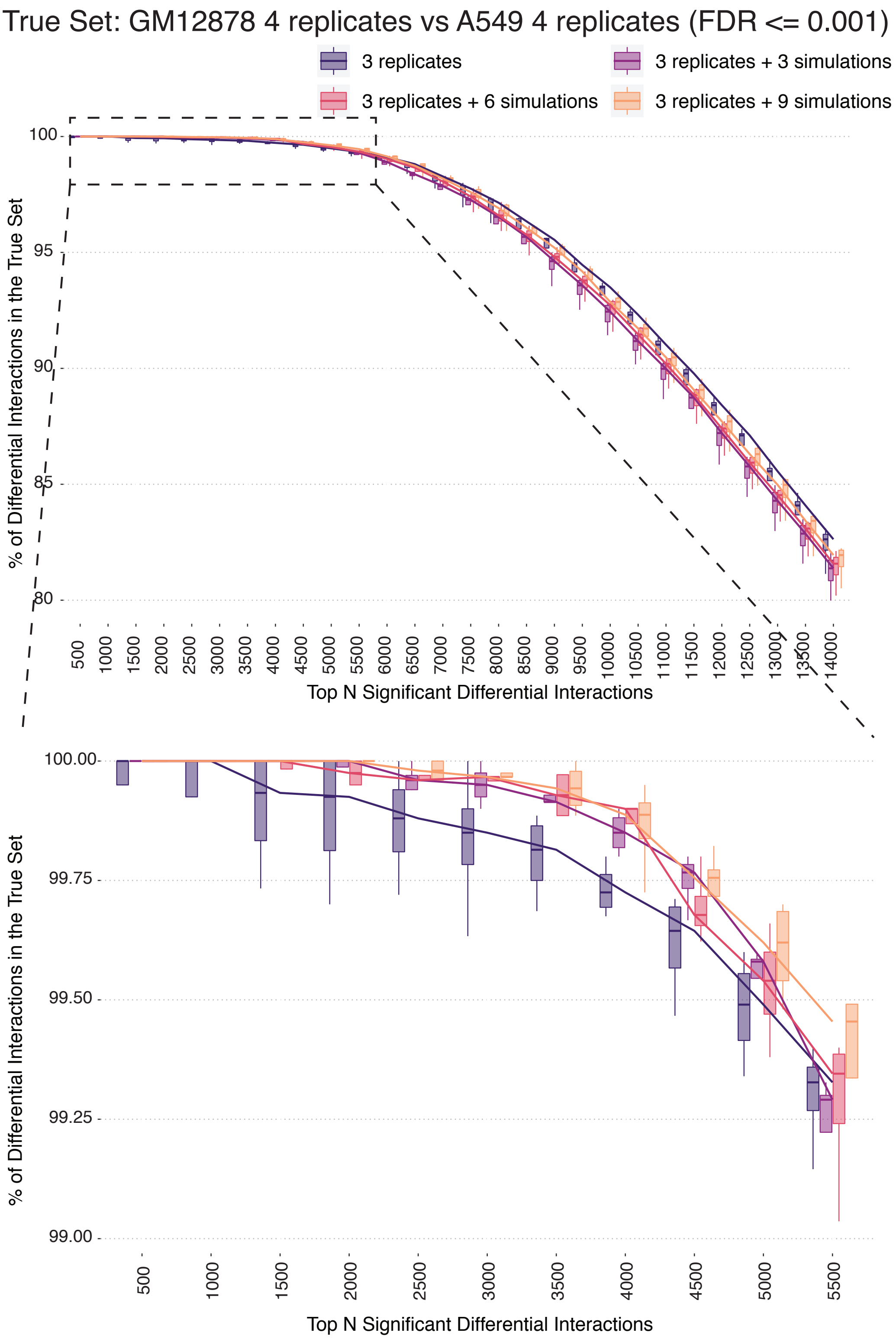


**Supplementary Figure 26 Percentage of top significant DCIs in the "true" differential chromatin interaction list, that is based on the comparison of the full set of 4 replicates of GM12878 with 4 replicates of A549, under the general setting of multiple replicates per condition and the corresponding FreeHi-C data augmentation settings.** Each condition includes 3 biological replicates. The "true" differential chromatin interaction set is defined as the most significant interactions in the comparison of 4 replicates of GM12878 with 4 replicates of A549 thresholded at

FDR of 0.001. Leveraging FreeHi-C replicates boosts the precision of top significant DCIs.

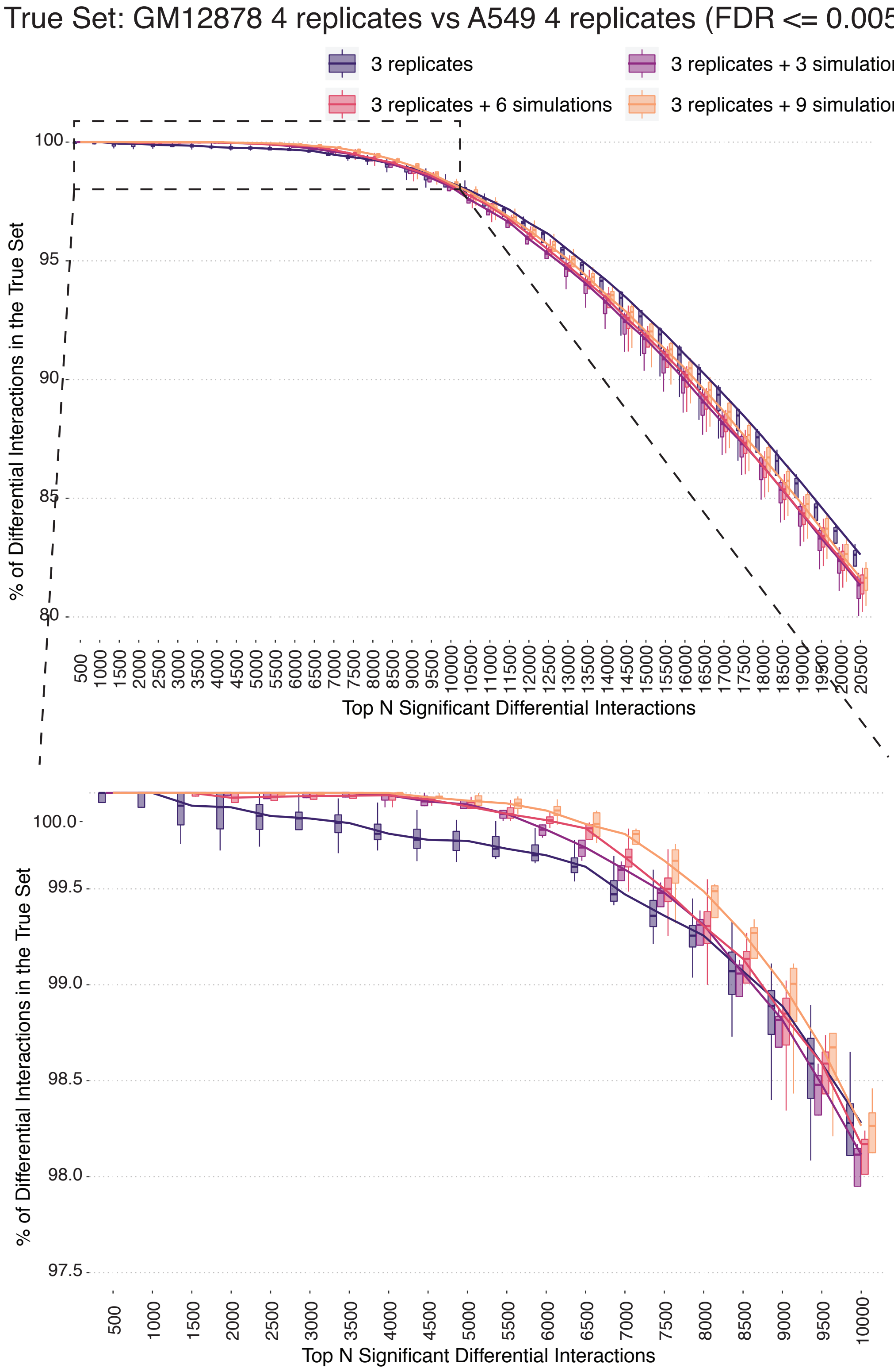


**Supplementary Figure 27 Same as Supplementary Fig. 26 at a different FDR threshold of 0.005 for defining the true set.**

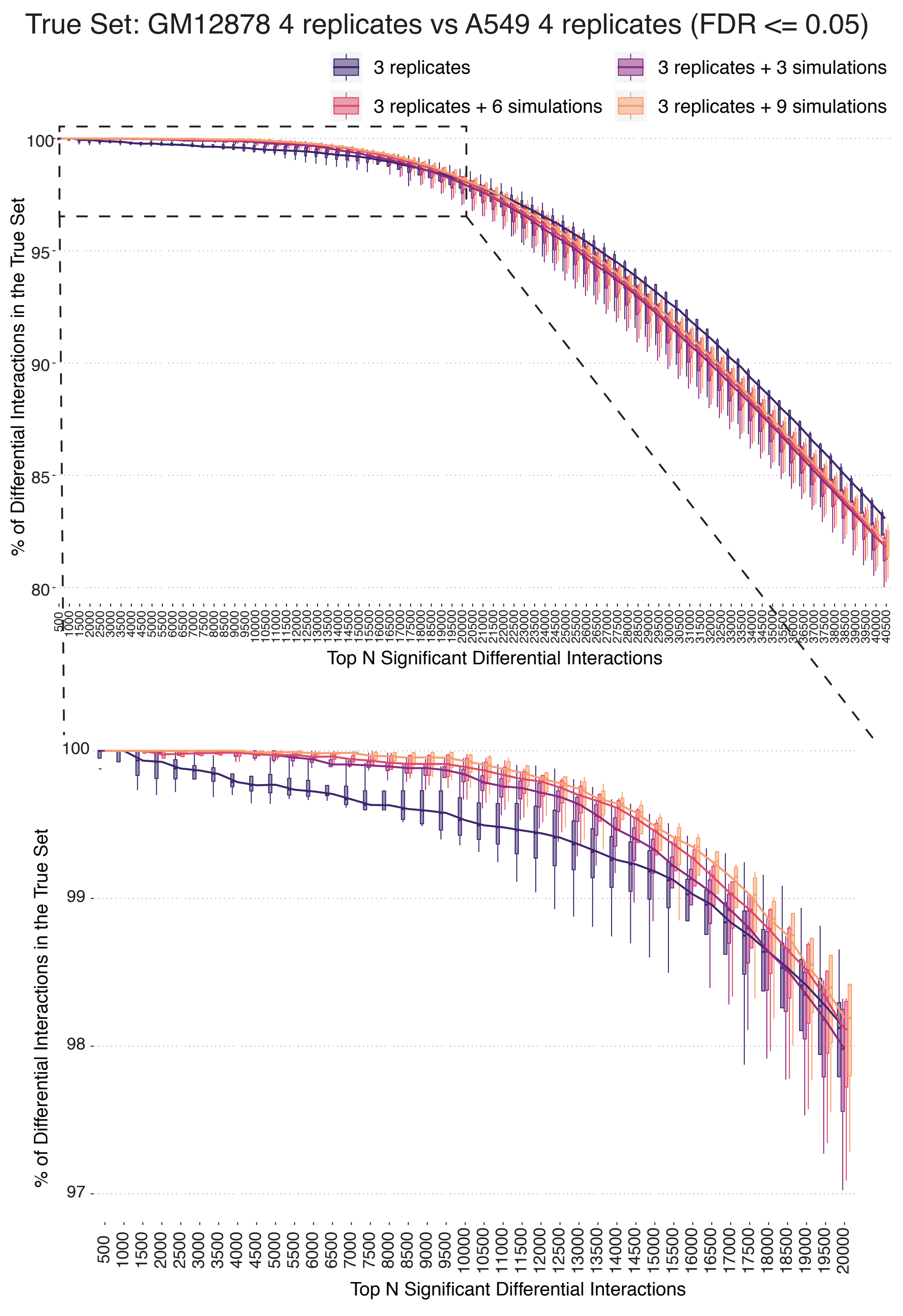


**Supplementary Figure 28 Same as Supplementary Fig. 26 at a different FDR threshold of 0.05 for defining the true set.**

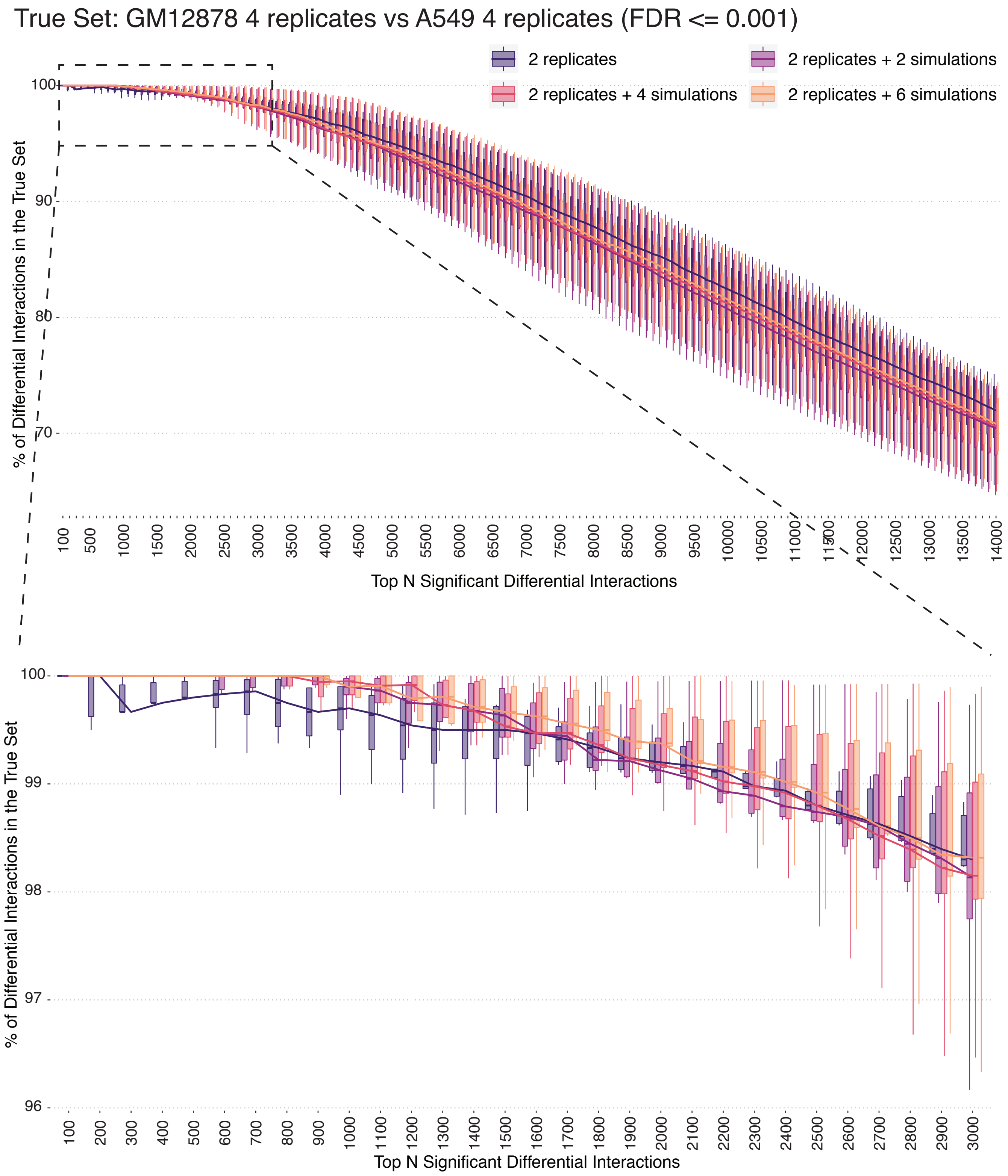


**Supplementary Figure 29 Percentage of top significant DCIs in the "true" differential chromatin interaction list, that is based on the comparison of full set of 4 replicates of GM12878 with 4 replicates of A549, under the general setting of multiple replicates per condition and the corresponding FreeHi-C data augmentation settings.** Each condition includes 2 biological replicates. The "true" differential chromatin interaction set is defined as the most significant interactions in the comparison of 4 replicates of GM12878 with 4 replicates of A549 thresholded at FDR of 0.001. Leveraging FreeHi-C replicates boosts the precision of top significant DCIs.

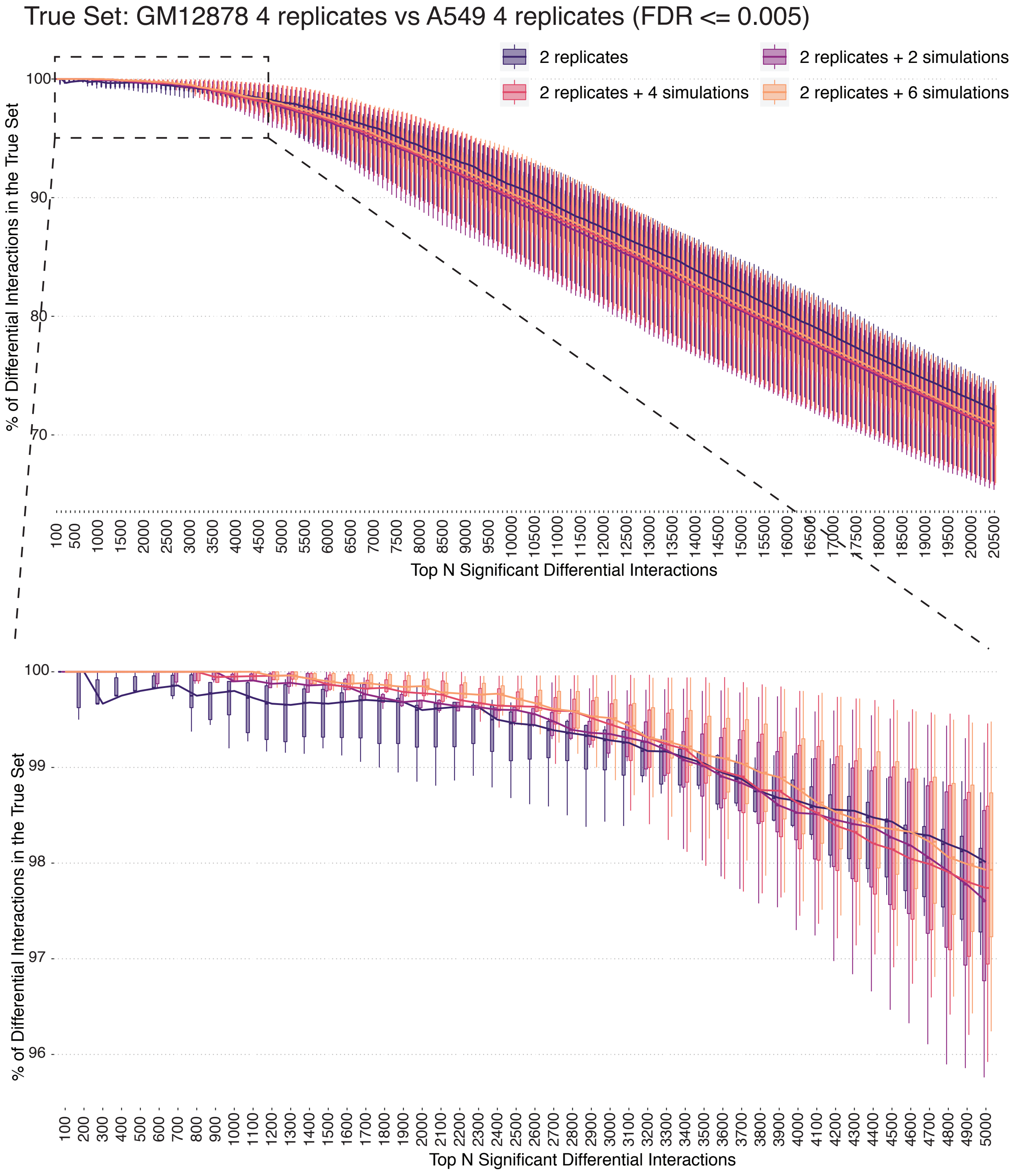


**Supplementary Figure 30 Same as Supplementary Fig. 29 at a different FDR threshold of 0.005 for defining the true set.**

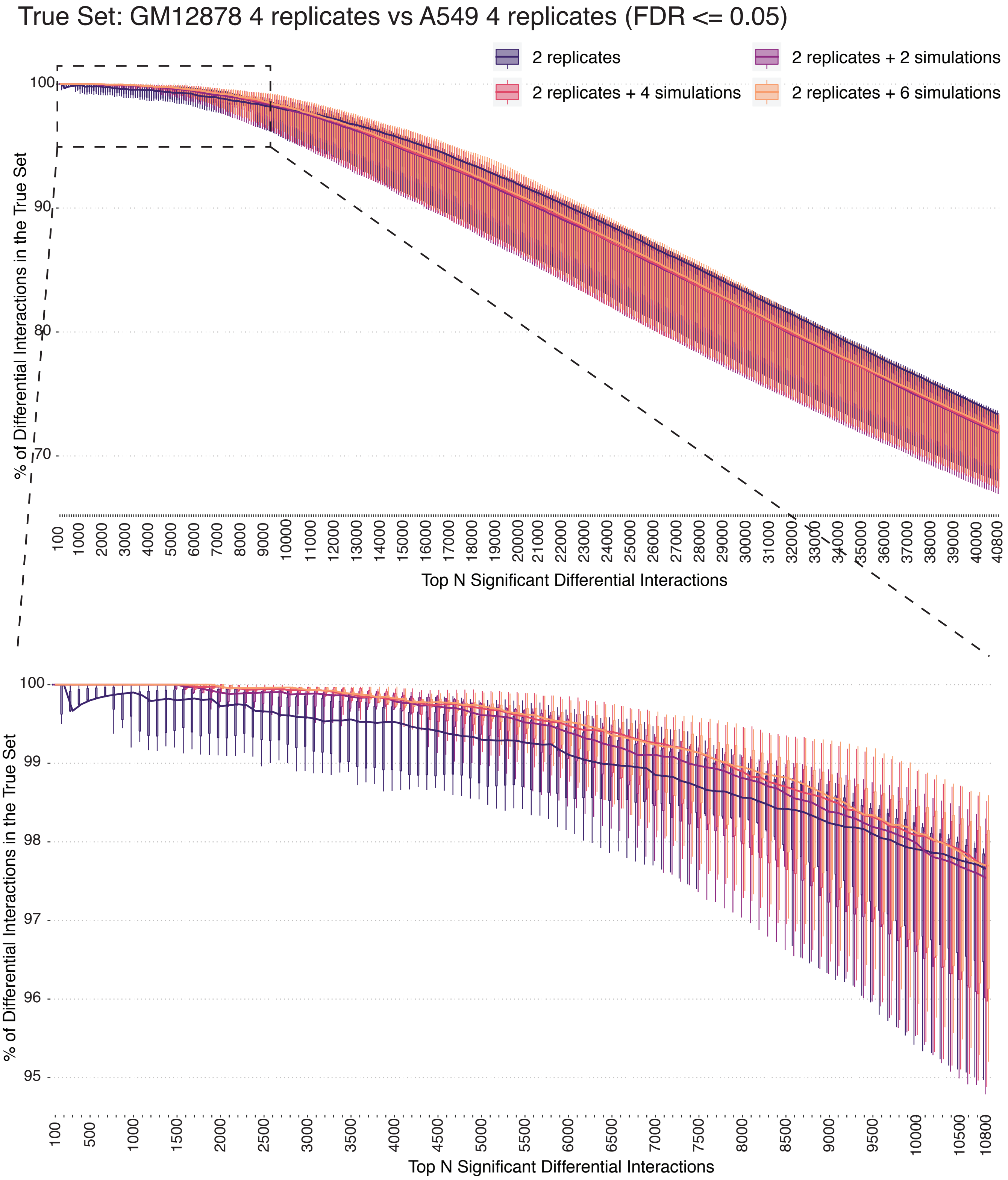


**Supplementary Figure 31 Same as Supplementary Fig. 29 at a different FDR threshold of 0.05 for defining the true set.**

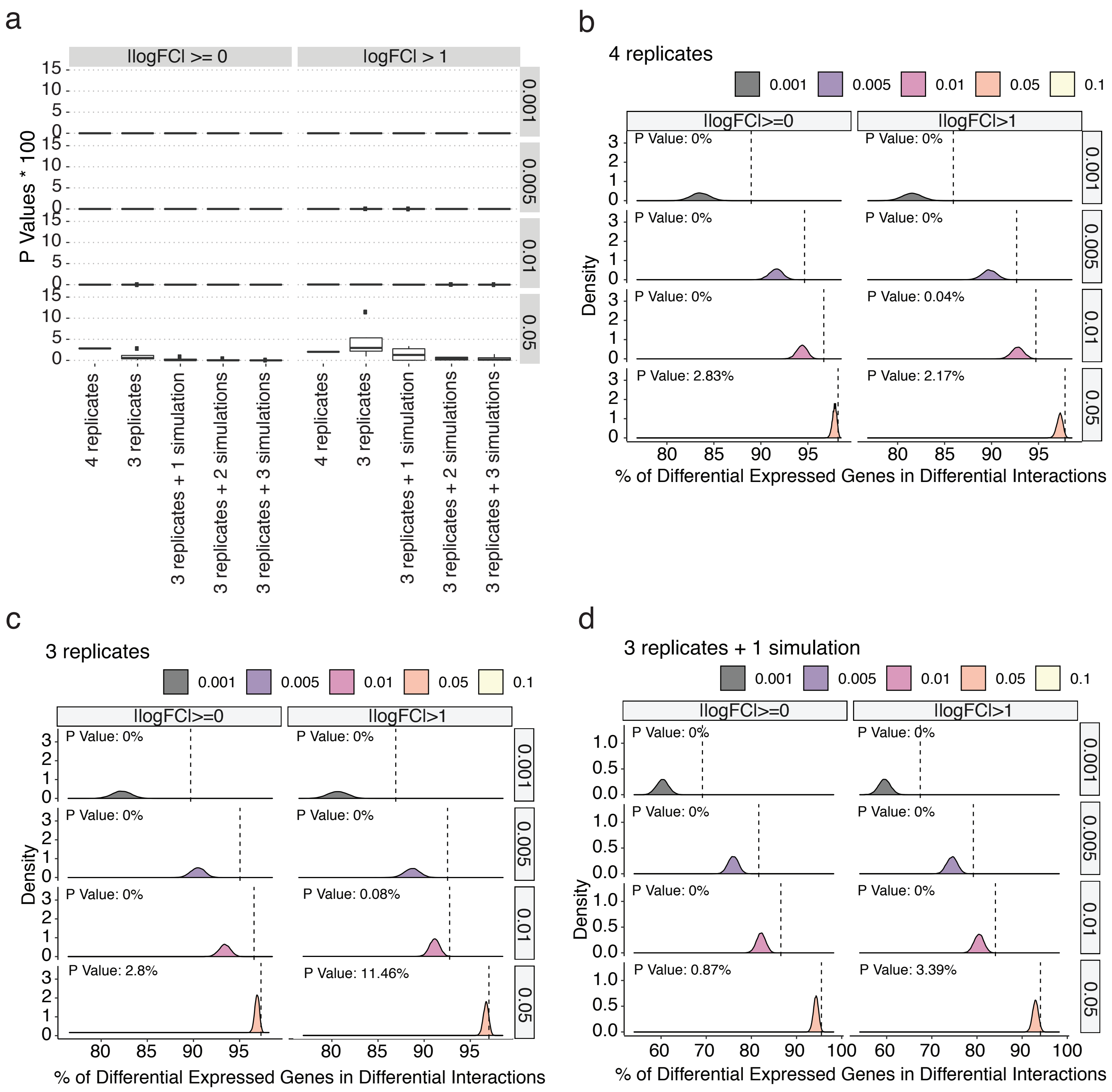


**Supplementary Figure 32 Evaluation of differential chromatin interactions detected by the three replicates per condition settings with external RNA-seq data of GM12878 and A549 cells.** **a** Significance of overlap of DCIs detected at varying FDRs with the differentially expressed genes between GM12878 and A459 cells. **b-d** are the individual randomization test results for the corresponding comparisons in **a**: 4 biological replicates (**b**), 3 biological replicates (**c**), and 3 biological replicates plus one FreeHi-C simulation for each of the biological replicates (**d**). The dashed lines represent the observed percentages of differentially expressed genes overlapping with differential interactions.

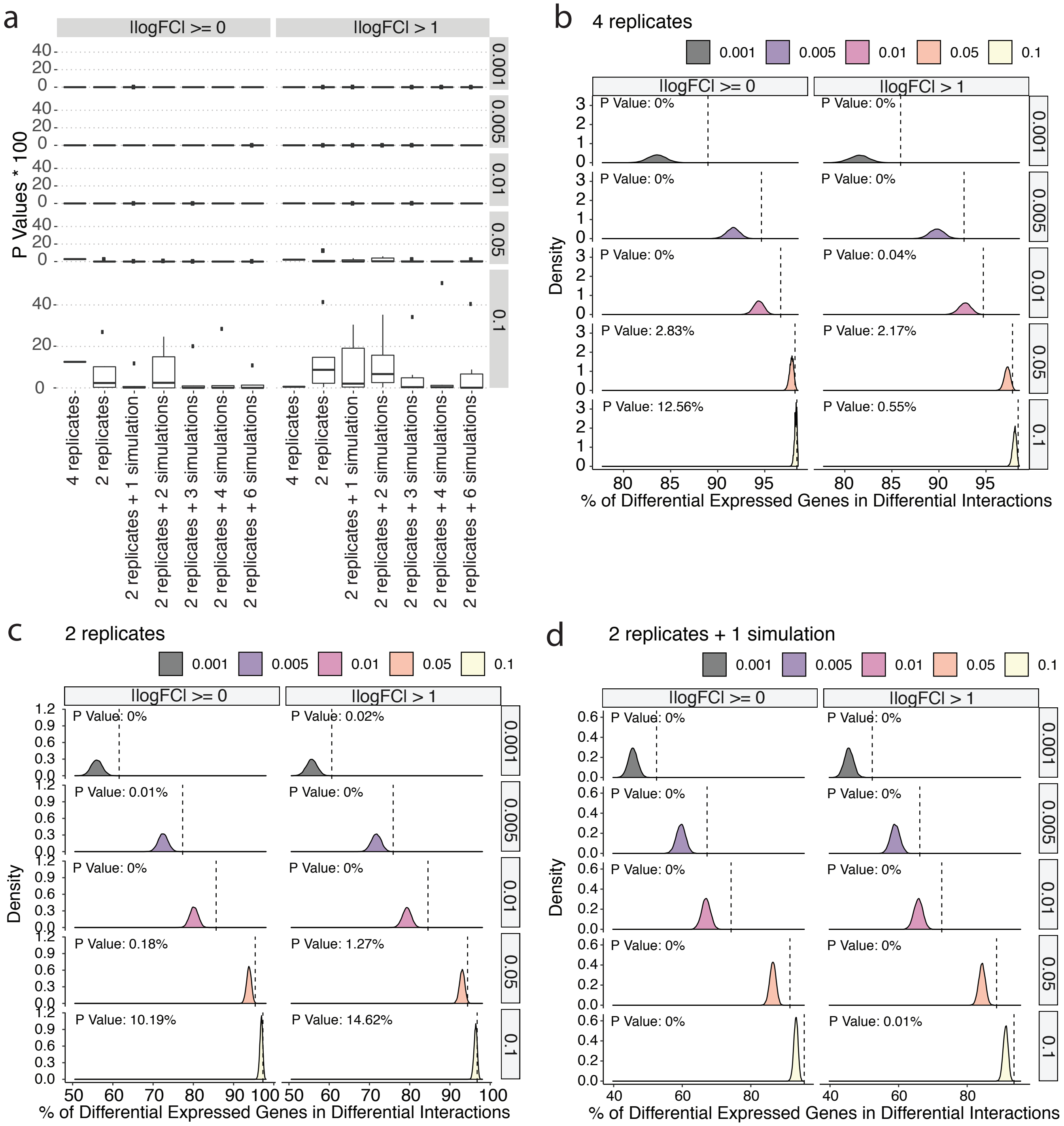


**Supplementary Figure 33 Evaluation of differential chromatin interactions detected by the two replicates per condition settings with external RNA-seq data of GM12878 and A549 cells. a** Significance of overlap of DCIs detected at varying FDRs with the differentially expressed genes between GM12878 and A459 cells. **b-d** are the individual randomization test results for the corresponding comparisons in **a**: 4 biological replicates (**b**), 2 biological replicates (**c**), and 2 biological replicates plus one FreeHi-C simulation for each of the biological replicates (**d**). The dashed lines represent the observed percentage of differentially expressed genes overlapping with differential interactions.

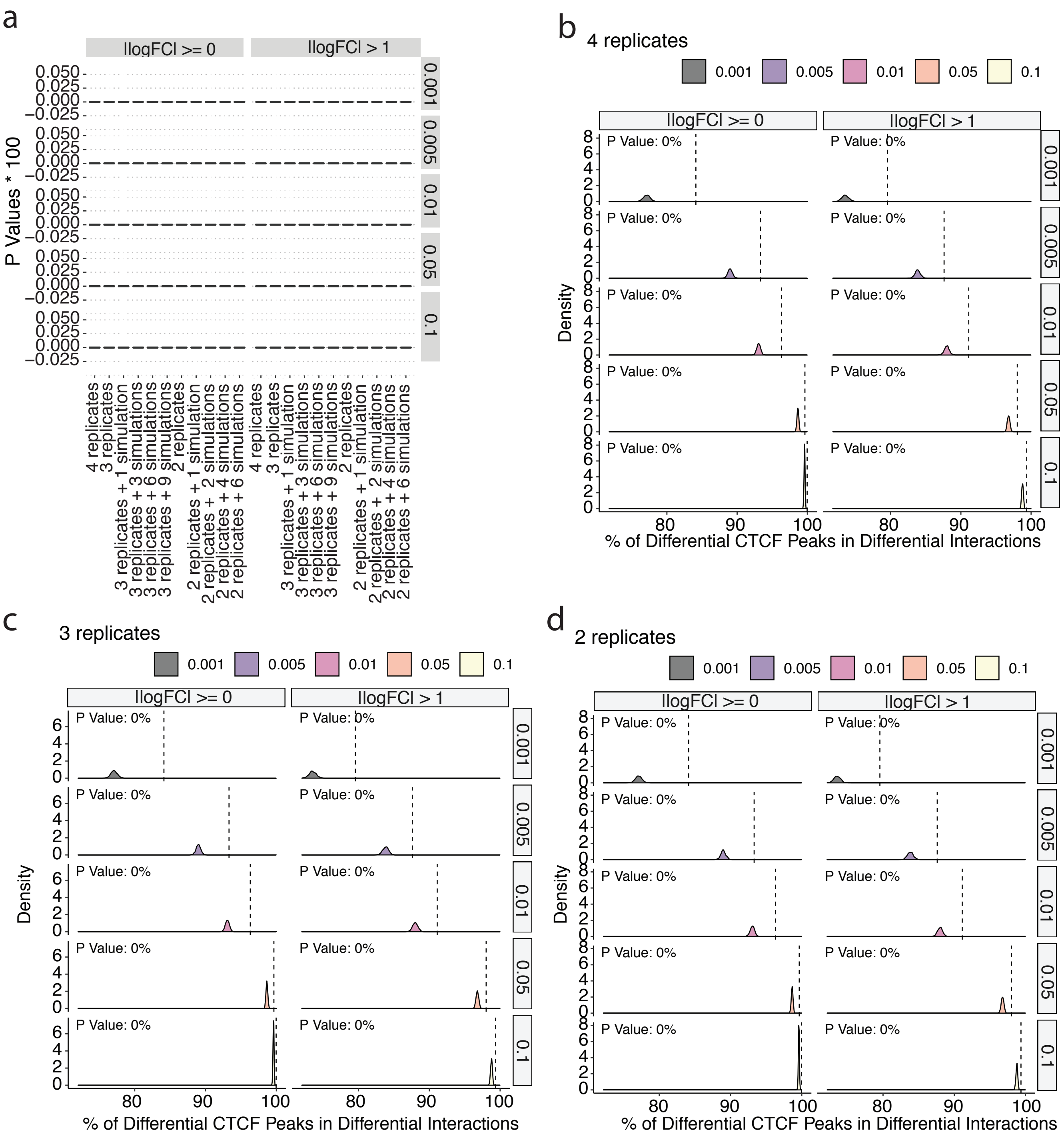


**Supplementary Figure 34 Evaluation of differential chromatin interactions detected with external differential CTCF ChIP-seq data of GM12878 and A549 cells.** **a** Significance of overlap of DCIs detected at varying FDRs with the CTCF ChIP-seq peaks. **b-d** are the individual randomization test results for the corresponding comparisons in **a**: 4 biological replicates (**b**), 3 biological replicates (**c**), and 2 biological replicates (**d**). The dashed lines represent the observed percentage of differential CTCF ChIP-seq peaks overlapping with differential interactions.

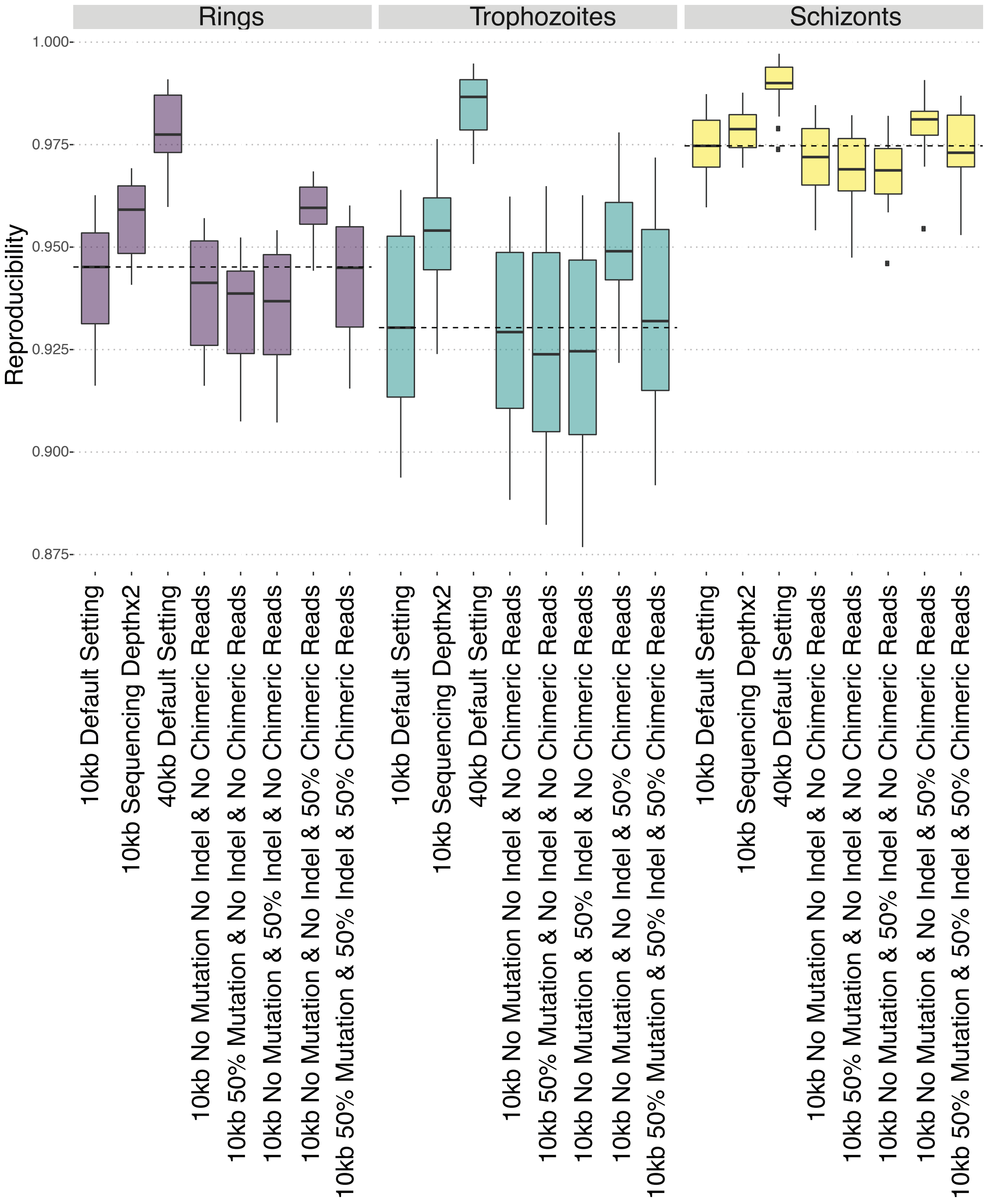


**Supplementary Figure 35 Evaluation of the effect of FreeHi-C parameters on the similarity between simulated replicates and the seed biological replicate using *P. falciparum*.** The reproducibility comparison evaluates the impact of resolution (10kb or 40kb), sequencing depth (original sequencing depth or 2 times the original sequencing depth), the proportion of mutation, indel, and chimeric reads.

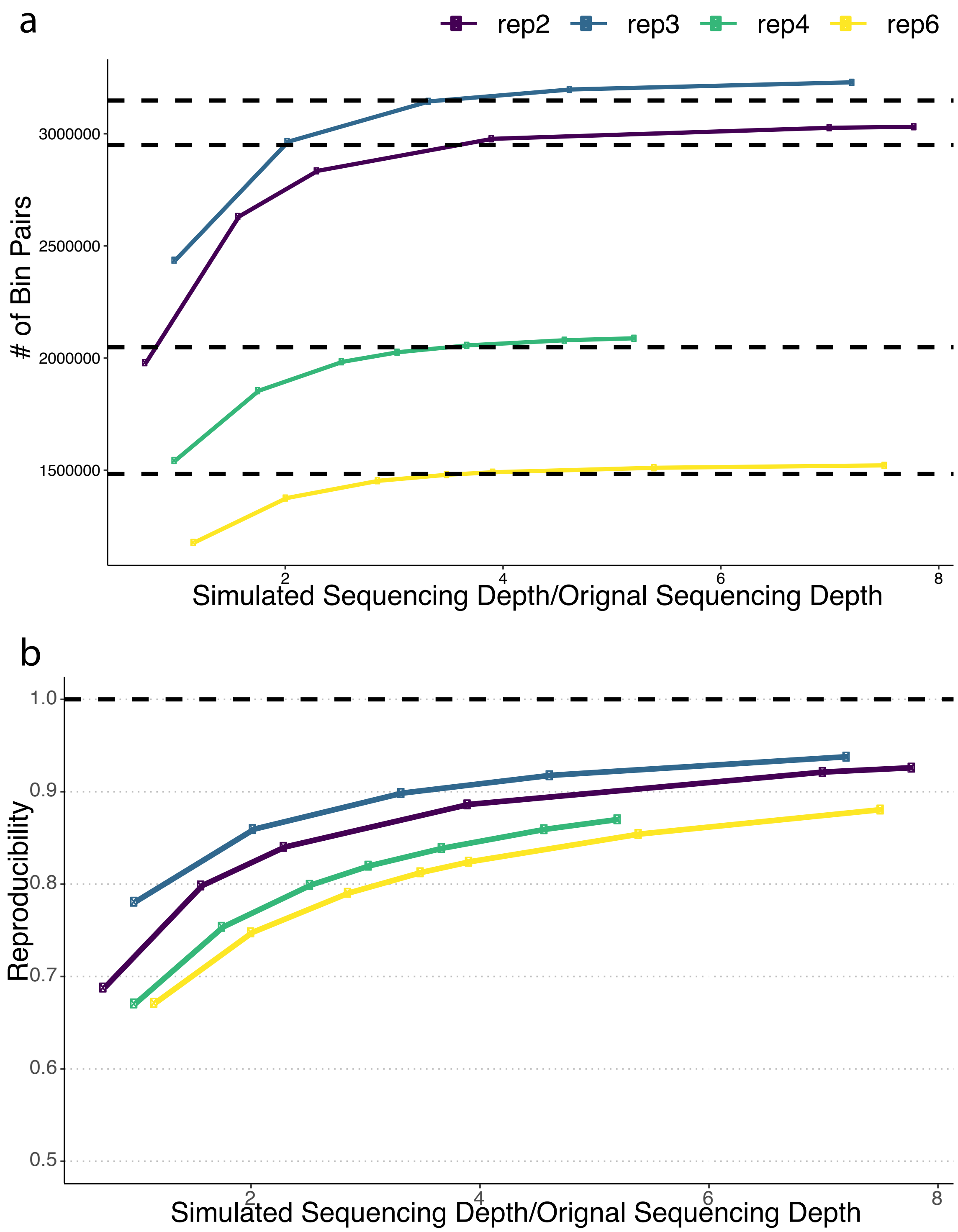


**Supplementary Figure 36 The impact of sequencing depth of simulation on the bin-pair coverage and HiCRep reproducibility with the seed biological replicates.** **a.** The numbers of unique bin-pair covered in the simulated replicates increase with the increasing sequencing depths. The dashed lines represent the bin-pair coverage in the four seed biological replicates. **b.** The HiCRep reproducibility between simulated replicates and seed biological replicates increases with the increasing simulated sequencing depth.

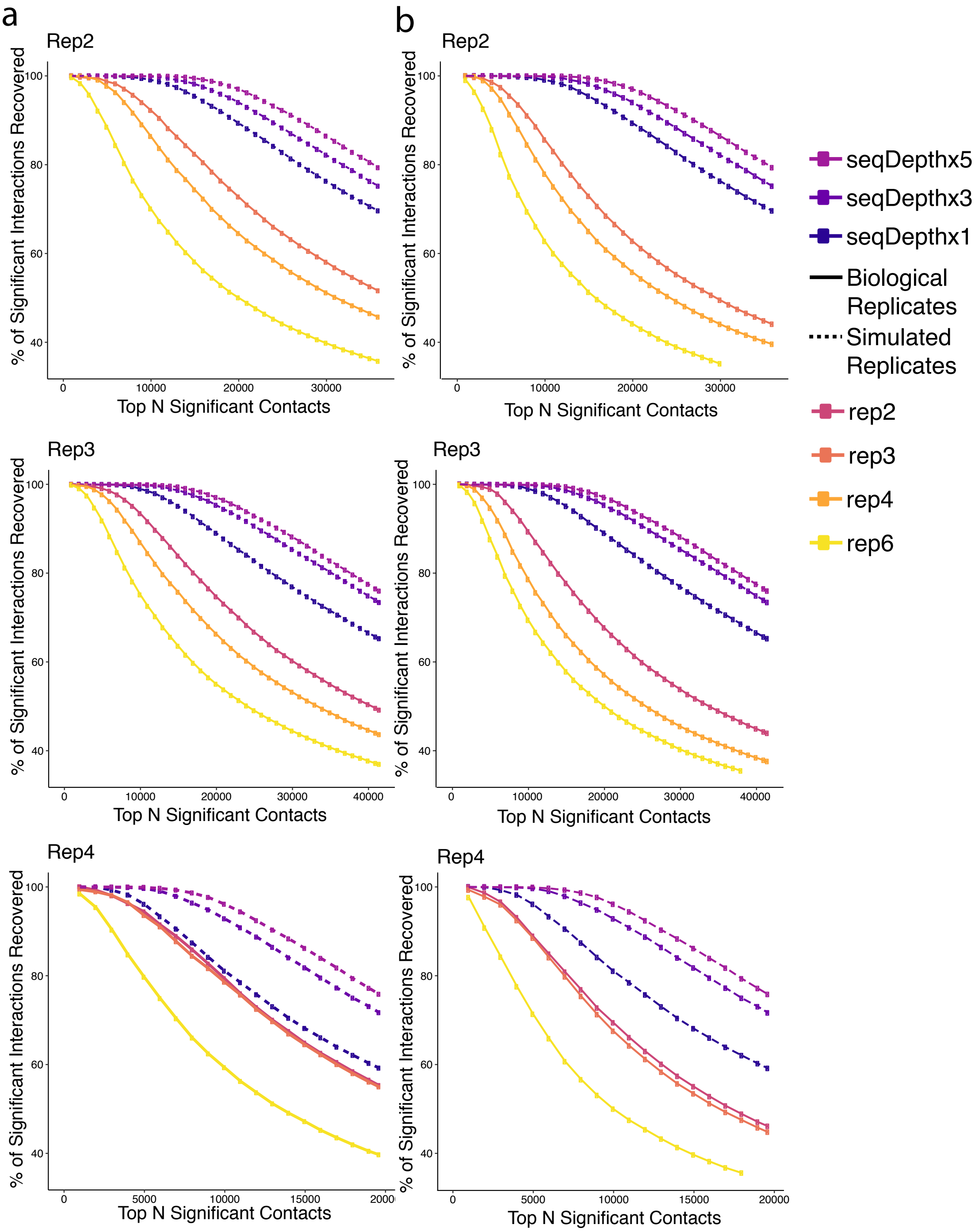


**Supplementary Figure 37 Percentage of the significant interactions in simulated replicates and other biological replicates of GM12878 that are also deemed significant in its original biological replicate.** The recovery rate of FreeHi-C simulated replicates of the targeted replicate at different sequencing depths are depicted in dashed purple, light blue and dark blue and are the same for **a** and **b** of the same targeted replicates. **a.** Biological replicates used in the significant interaction detection are the original (seed) replicates. **b.** Biological replicates (i.e., not the targeted biological replicate) used in significant interaction detection are the replicates simulated to the sequencing depth of the targeted biological replicate listed on the top of each figure panel.

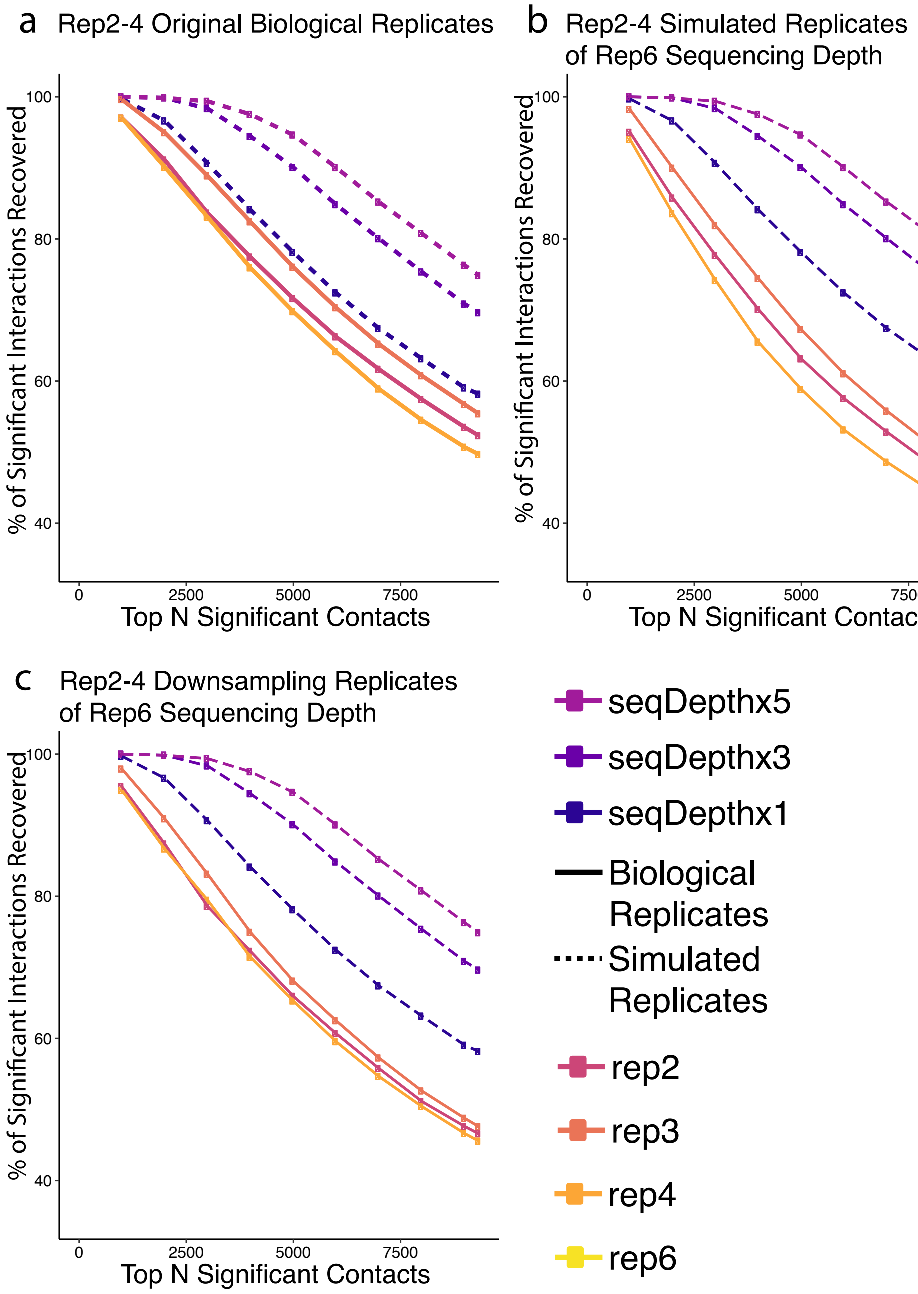


**Supplementary Figure 38 Percentage of the significant interactions in simulated replicates and other biological replicates of GM12878 that are also deemed significant in its original biological replicate6.** The targeted biological replicate is replicate6 of GM12878. The recovery rate of FreeHi-C simulated replicates of replicate6 at different sequencing depths are depicted in dashed purple, light blue and dark blue and are the same across **a-c**. **a.** Biological replicates used in significant interaction detection are the original (seed) replicates. **b.** Biological replicates (i.e., not the targeted biological replicate) used in significant interaction calling are the replicates simulated to the sequencing depth of the targeted biological replicate6. **c.** Biological replicates (i.e., not the targeted biological replicate) used in significant interaction calling are the replicates downsampled to the sequencing depth of the targeted biological replicate6.
